# Cytoplasmic Abundant Heat-Soluble Proteins from Tardigrades Protect Synthetic Cells Under Stress

**DOI:** 10.1101/2025.08.07.669088

**Authors:** Yongkang Xi, Jianming Mao, Samuel J. Chen, Hossein Moghimianavval, Young Jin Lee, Ayush Panda, Alexander J. Huang, Daniel H. Zhou, L. Andy Xu, Kayla Y. Fu, Solomon Adera, Andrew L. Ferguson, Allen P. Liu

**Affiliations:** Department of Mechanical Engineering, University of Michigan, Ann Arbor, MI, 48109 USA; Department of Chemistry, University of Chicago, Chicago, IL, 60637; Department of Biophysics, University of Michigan, Ann Arbor, MI, 48109 USA; Pritzker School of Molecular Engineering, University of Chicago, Chicago, IL, 60637; Department of Biomedical Engineering, University of Michigan, Ann Arbor, MI, 48109 USA; Cellular and Molecular Biology Program, University of Michigan, Ann Arbor, MI, 48109 USA

## Abstract

Cytoplasmic abundant heat-soluble (CAHS) proteins, a stress-responsive intrinsically disordered protein from tardigrades, have been discovered to form gel-like networks providing structural support during dehydration, thus enabling anhydrobiosis. However, the mechanism by which CAHS proteins protect the dehydrating cellular membrane remains enigmatic. Using giant unilamellar vesicles (GUVs) as a model membrane system, we show that encapsulated CAHS12 undergoes a reversible structural transformation that reinforces membrane integrity and preserves encapsulated components, mimicking natural anhydrobiosis. CAHS12-containing GUVs demonstrated stability for weeks and mechanical robustness under dehydration, elevated temperature, and osmotic stresses. Molecular simulations suggest that CAHS12 forms a filamentous network within the vesicle lumen that mitigates membrane collapse and preserves compartmental architecture. Synthetic cells with cell-free transcription-translation capabilities withstand desiccation and recover biochemical activities, akin to the tun state of the tardigrade. This discovery opens up synthetic cell applications in bioengineering, cold-chain-independent biomanufacturing, and adaptive biointerfaces.

---

Nature provides compelling examples of organisms that remain viable in near-total absence of water^1^. Tardigrades exemplify this extreme resilience^2,3^. These microscopic invertebrates survive complete desiccation, vacuum, ionizing radiation, and extreme temperatures by entering an ametabolic state known as anhydrobiosis, during which cellular water is expelled, and biological activity is halted indefinitely^4,5^. Remarkably, they avoid molecular damage—such as protein denaturation, membrane collapse, and functional impairment—commonly associated with desiccation. While classical models emphasize osmolytes like trehalose or antioxidants^6^, growing evidence implicates a family of tardigrade intrinsically disordered proteins, namely the cytoplasmic-abundant heat-soluble (CAHS) proteins, as the primary drivers of this exceptional dry-state tolerance^7^.

Tardigrades possess a diverse repertoire of cytoplasmic abundant heat-soluble (CAHS) proteins with partially overlapping yet isoform-specific functions under stress. Under hydrated conditions, CAHS proteins are largely unstructured; upon dehydration or osmotic stress, they undergo reversible, stress-responsive conformational transitions, with intrinsically disordered N-and C-terminal regions flanking a central α-helical linker domain (Fig. 1A). Early biochemical and biophysical studies demonstrated that CAHS proteins can vitrify and suppress protein aggregation under water loss, implicating them as molecular stabilizers during anhydrobiosis^2,8^. Previous studies showed that CAHS proteins form gel-like or filamentous assemblies *in vitro* and in cells under dehydration or crowding, and that these assemblies correlate with enhanced mechanical resilience and cellular survival (**Supplementary Fig. 1**)^9–11^.

**Fig. 1.**
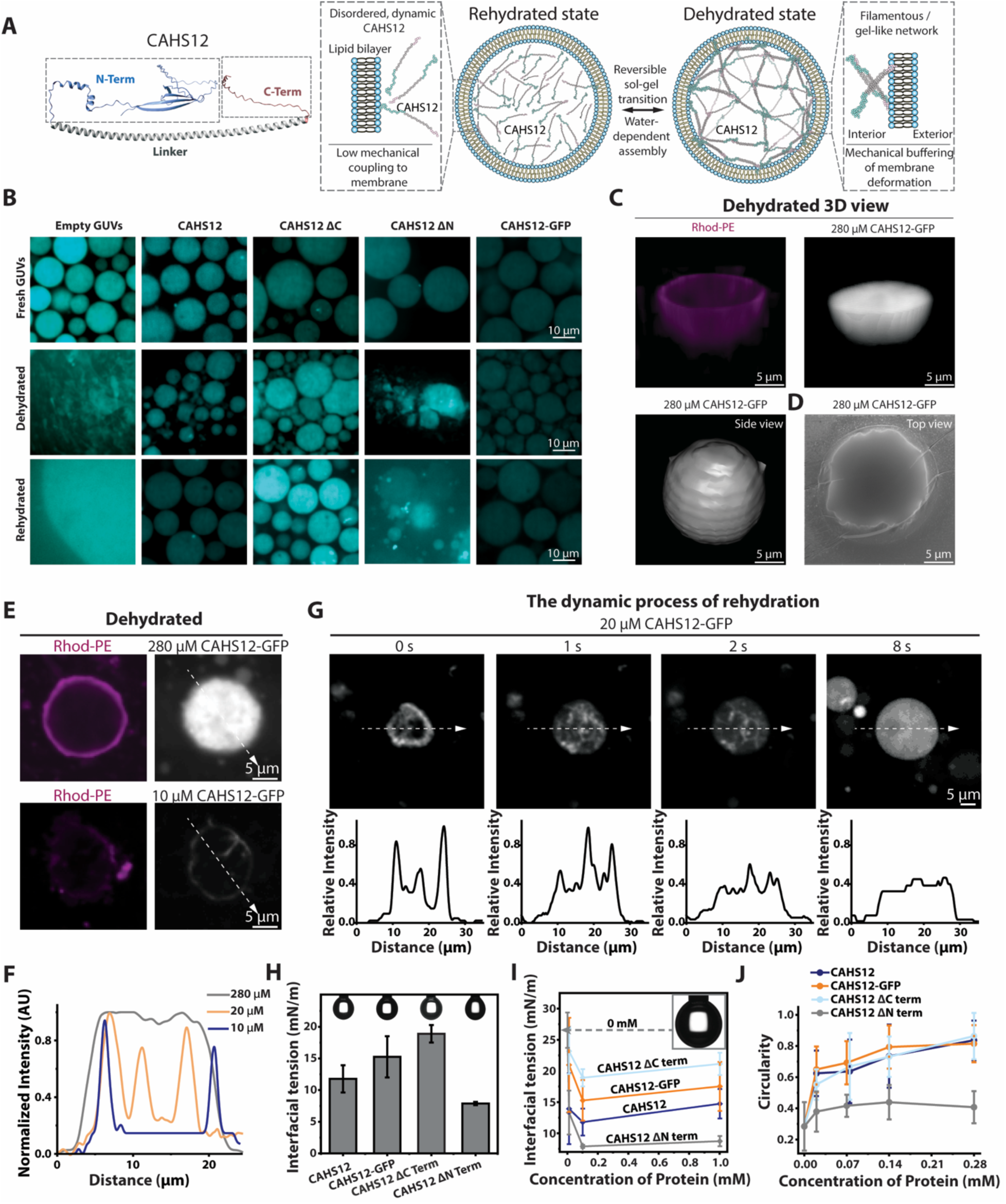
Structural configuration and functional role of CAHS12 in desiccation-tolerant GUVs. (**A**) Proposed model of CAHS12-mediated stabilization of GUV membranes during dehydration and rehydration. (left) Structural organization of CAHS12, showing an intrinsically disordered N-terminal region (blue), a central α-helical linker (gray), and a disordered C-terminal region (red); (right) Schematic illustrating CAHS12 behavior during rehydration: CAHS12 remains highly disordered and dynamically distributed within GUVs, exhibiting transient homotypic interactions. Upon dehydration, reduced water activity induces reversible CAHS12 self-association into an extended filamentous or gel-like network within the vesicle lumen. This network mechanically couples to the confined volume and buffers membrane deformation without requiring stable membrane insertion. Rehydration restores the disordered, sol-like state. This model is consistent with prior reports of dehydration-induced CAHS gelation and filament formation. Molecular rendering of the CAHS12 protein was constructed using ChimeraX^81^. (**B**) Fluorescence microscopy images of CAHS12-loaded GUVs under fresh, dehydrated, and rehydrated states. Cy5 encapsulated within GUV lumen was used as a leakage reporter to assess membrane integrity based on its release. The CAHS12 domain architecture shown is based on previous studies, and the experiments here examine how these domains contribute to intraluminal assembly and protection of GUV membranes. Z-axis scan 3D fluorescence microscopy image (**C**) and scanning electron microscopy image (**D**) of dehydrated GUVs. Magenta indicates membrane-localized Rhod-PE, while CAHS12-GFP is shown in gray (right and bottom left). (**E**) Fluorescence microscopy image of dehydrated GUVs containing 280 μM or 10 μM of CAHS12-GFP. (**F**) Line scan analysis of normalized GFP fluorescence across dehydrated GUV containing 10, 20, or 280 µM CAHS12-GFP; Scale bar, 5 µm. (**G**) The dynamic process of dehydration and rehydration of GUVs containing 20 μM of CAHS12-GFP; Scale bar, 5 µm. (**H**) Quantification of IFT of droplets containing 0.1 mM of various CAHS12 constructs in mineral oil (with 0.41 mM POPC). An example of drop shape is shown for each condition. (**I**) IFT as a function of concentration for CAHS12, CAHS12 ΔC, and CAHS12 ΔN. Inset shows the morphology of a pure water droplet suspended in mineral oil containing 0.41 mM POPC. (**J**) Circularity analysis of dehydrated GUVs with increasing concentrations of different CAHS12 constructs (n > 100). Error bars show standard deviation.

High-resolution structural and spectroscopic studies further refined this view by revealing that CAHS proteins remain highly dynamic and only partially ordered even within stress-induced assemblies. In particular, NMR and scattering analyses from the Blackledge laboratory demonstrated that CAHS proteins form adaptable, weakly interacting networks rather than rigid scaffolds, supporting a model of emergent protection driven by multivalent, reversible interactions^12^. More recently, cellular imaging and genetic perturbation studies showed that CAHS proteins reversibly reorganize into filamentous networks during dehydration and rapidly dissolve upon rehydration without compromising cell viability, reinforcing their role as stress-responsive biomolecular materials rather than static structural elements^11^.

Despite these advances, prior studies have largely focused on molecular assembly, material properties, or organismal survival. Whether CAHS proteins can preserve functional biochemical activity, particularly over extended periods of dry storage, has not been directly tested. Moreover, most demonstrations to date have been conducted in living cells or complex extracts, where endogenous repair pathways, chaperones, and metabolic processes obscure the individual contributions of CAHS-mediated membrane stabilization and protein protection.

Here, we focus on CAHS12, a conserved and abundantly expressed CAHS isoform induced during desiccation, previously shown to form reversible filamentous assemblies under stress^9^. To isolate the mechanism from cellular complexity, we reconstituted CAHS12 and its truncation variants within giant unilamellar vesicles (GUVs) as a minimal synthetic cell platform. This reductionist system enables direct interrogation of CAHS-mediated membrane stabilization and long-term functional preservation under dehydration. Notably, we demonstrate that CAHS12 enables sustained preservation of cell-free transcription–translation activity and membrane protein function within dehydrated GUVs, extending prior structural and cellular observations to long-term functional protection in synthetic cells.

## Results and discussions

### CAHS12 enables robust structural preservation of synthetic cells during dehydration-rehydration cycles

To determine whether cytoplasmic proteins from tardigrades can confer desiccation tolerance to GUVs, we encapsulated a panel of candidate proteins and subjected them to dehydration–rehydration cycles (**Fig. 1B, Supplementary Fig. 2-3**). GUVs encapsulating *Ramazzottius varieornatus* CAHS12-GFP maintained their size after drying and rehydration (8.4 ± 4.1 μm before drying vs. 7.0 ± 2.3 μm after rehydration; n > 100, **Supplementary Fig. 3C**). In contrast, GUVs containing bovine serum albumin (BSA), actin, GFP or no protein underwent aggregation, collapse, or rupture during drying and exhibited only partial recovery. These failures likely reflect membrane stresses arising from steep osmotic gradients and evaporative flux, which compress the bilayer, thin the hydration shell, enhance van der Waals adhesion, and destabilize the air–water interface ^13–15^. Without a stabilizing mechanism, these forces lead to irreversible structural failure and cargo leakage (*e.g.*, Cy5) upon rehydration.

To identify the structural determinants of CAHS12-mediated protection, we generated fluorescent fusion proteins and truncation variants removing either the C or N terminus, hereafter referred to as ΔC and ΔN (**Supplementary Fig. 2 and 15**, **Supplementary Table 1, 4, 5**). GUVs encapsulating full-length CAHS12, -GFP or -BFP retained spherical morphology even at 20 μM, indicating that the bulky fluorescent tags do not impair function (**Fig. 1B**, **Supplementary Fig. 4-5 and 6A and Supplementary Table 7**). Likewise, the ΔC variant preserved vesicle shape, suggesting the C-terminal domain is unnecessary (**Supplementary Fig. 6B**). By contrast, the ΔN variant failed to prevent GUV collapse or cargo leakage, even at 280 μM, indicating a critical role for the N-terminal intrinsically disordered regions in protective function (**Supplementary Fig. 6C**). To investigate how CAHS12 preserves vesicle architecture, we imaged dried GUVs using confocal and scanning electron microscopy. Three-dimensional reconstructions showed that GUVs containing 280 μM CAHS12-GFP formed compact but intact spherical structures, with continuous membranes labeled by rhodamine-PE lipids (**Fig. 1C**). GFP fluorescence was evenly distributed across both the membrane and inside. Scanning electron microscopy confirmed smooth, rounded vesicle morphology (**Fig. 1D**), consistent with resistance to interfacial compression and membrane rupture. To probe concentration dependence, we compared CAHS12-GFP at 280, 20, and 10 μM. At high concentration (i.e., 280 μM), CAHS12-GFP appeared uniform (**Fig. 1E, 1F**). At lower concentrations, CAHS12-GFP was distributed to the membrane interface, consistent with cooperative membrane binding via multivalent interactions or weak amphiphilic motifs within the disordered regions (**Fig. 1F, 1G**). Real-time imaging during rehydration revealed rapid morphological recovery (**Fig. 1G**). GUVs with 20 μM CAHS12-GFP expanded within 1 s of water addition and regained spherical geometry by 8 s, while GFP signal redistributed from uneven to uniform across the membrane, reflecting dynamic re-equilibration with decreasing membrane tension. Similar results were observed during the rehydration of GUVs containing 10 μM CAHS12-GFP (**Supplementary Fig. 7**). These observations suggest that CAHS12-GFP forms a reversible, stress-responsive network that enables compression tolerance and prevents membrane rupture and content loss during desiccation–rehydration transitions.

To understand the physical basis for this protection, we assessed CAHS12’s interfacial properties using pendant drop tensiometry^16^. In the presence of POPC (0.41 mM), CAHS12 lowered interfacial tension (IFT) from ∼26 to ∼12 mN/m at 0.1 mM, indicative of surface adsorption and tension buffering (**Fig. 1H**)^17,18^. To further quantify this interfacial adsorption behavior, we estimated the apparent surface excess concentration (Γ) using the Gibbs adsorption equation, 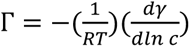, where Γ is the surface excess concentration (mol/m^2^), γ is the interfacial tension (N/m), *c* is the bulk protein concentration (mol/L), *R* is the universal gas constant (8.314 J·mol^-1^·K^-1^), and *T* is the absolute temperature (K). Over the concentration range of 0.1–1.0 mM, CAHS12 exhibited a modest increase in interfacial tension (from ∼12 to ∼15 mN/m), corresponding to a surface excess of approximately 5.3×10^−7^ mol/m^2^. Although the ΔN variant exhibited a smaller absolute increase in interfacial tension (from ∼6 to ∼8 mN/m), the steeper slope in γ vs. ln *c* yielded a larger estimated surface excess (Γ ≈ 1.4×10⁻⁶ mol/m²), suggesting more extensive interfacial accumulation. It may not form a cohesive or stress-buffering network—consistent with its inability to protect vesicle morphology or prevent cargo leakage upon rehydration. This disparity between adsorption quantity and protective efficacy implies that interfacial coverage alone is insufficient; rather, the structural organization and molecular interactions facilitated by the N-terminal domain are likely critical for forming a functional interfacial scaffold. The non-monotonic behavior of CAHS12 (i.e., IFT increasing at higher concentrations) further hints at possible crowding or structural rearrangements at saturated interfaces, which may reflect dynamic assembly processes unique to the full-length protein. At 1 mM, IFT increased slightly (∼15 mN/m), possibly due to interfacial saturation or crowding (**Fig. 1I**)^19^. Both CAHS12-GFP and ΔC variants retained this activity, while ΔN constructs showed minimal IFT reduction (<10 mN/m), confirming the N-terminal region mediates lipid-specific surface interaction. In the absence of POPC, all constructs showed similarly IFT (∼20–28 mN/m), indicating that although CAHS12 possesses intrinsic surface activity, its ability to regulate interfacial tension is strongly enhanced at lipid interfaces (**Supplementary Fig. 8**). Consistent with these interfacial measurements, we performed micropipette aspiration experiments of CAHS12-containing GUVs to further demonstrate construct- and concentration-dependent mechanical stabilization of GUVs by CAHS12 (**Supplementary Fig. 9**). Increasing concentrations of CAHS12–GFP (0.05 mM to 0.90 mM) confer progressively greater resistance to deformation under applied membrane tension, while comparison of truncation variants reveals that full-length CAHS12 and the ΔC construct effectively stabilize GUVs, whereas the ΔN variant fails to do so and the GUV undergoes pronounced elongation during aspiration. We next investigated whether these interfacial properties underlie membrane stabilization. Circularity analysis of dried GUVs showed that full-length and CAHS12 ΔC maintained shape integrity (circularity > 0.85), whereas the ΔN variant yielded deformed vesicles (circularity < 0.4), consistent with structural failure (**Fig. 1J** and **Supplementary Fig. 10**). These results demonstrate that CAHS12 operates as a lipid-specific, disordered interfacial modulator that assembles reversibly at stressed membranes, buffers tension, and enables rapid vesicle recovery following rehydration.

### Molecular dynamics simulations of CAHS12 protein assemblies reveal key interactions with phospholipid membranes

To probe the underlying mechanism by which CAHS12 protects GUVs during dehydration, we employed coarse-grained molecular dynamics (MD) simulations using the Martini3 force field^20^ to model the protein dynamics, assembly behaviors, and interaction with a POPC lipid membrane (**Fig. 2A).** Inspired by prior studies^12,21,22^, the CAHS12 protein was modeled as a dumbbell-like construct where the N-terminal (residues 1-153) and C-terminal regions (residues 268-297) were represented as random coils and the linker region (residues 154-267) as an extended α-helix. This structural representation is supported by an analysis of the *κ* value as a measure of charge patterning within a protein sequence, with high values indicative of disordered conformations^23^: high local *κ* values in the terminal regions suggest disordered conformations, and low *κ* values in the linker region suggest ordered conformations. These secondary structure assignations result in a molecular model of a single CAHS12 protein in water possessing a radius of gyration (*R_g_*) of (5.21 ± 0.11) nm, which falls in a similar regime for other tardigrade disordered proteins from prior studies (**Supplementary Fig. 11**)^12,22,24^. Using this model, we first explored inter-protein interactions by simulating systems of two CAHS12 molecules. Principal components analysis (PCA)^25^ of the resulting dimer conformations reveals that CAHS12 dimers associate primarily through interactions of the C- and N-terminal regions (**Fig. 2B** and **Supplementary Fig. 12**). Notably, the N-terminal region mediates stronger binding with interaction energy (IE) averaging ∼(−280) kcal/mol, compared to ∼(−69) kcal/mol for C-terminal-only interactions, as measured by the short-range inter-protein interaction energy. We note that antiparallel CAHS12 dimer conformations have been predicted in prior studies using structure prediction tools^10^. Free energy calculations indicate that the antiparallel dimer is approximately 20 kcal/mol more stable than the parallel dimer (**Supplementary Fig. 13**), but we observe only the parallel dimer state in our unbiased calculations and no significant population of the antiparallel dimer. This suggests that while the antiparallel dimer may be more thermodynamically stable, it may be less kinetically accessible than the parallel dimer state. Furthermore, a large proportion of the observed dimers do interact via binding between the C- and N-terminal regions that may be precursors to antiparallel dimers in longer simulations^9,26,27^. In simulations of CAHS12 proteins at varying concentrations, protein assembly mediated by the N- and C-terminal regions showed a concentration-dependent trend, with higher concentrations leading to increased terminal close contacts per protein (**Supplementary Fig. 14**), a trend also observed in experiments^28^.

**Fig. 2.**
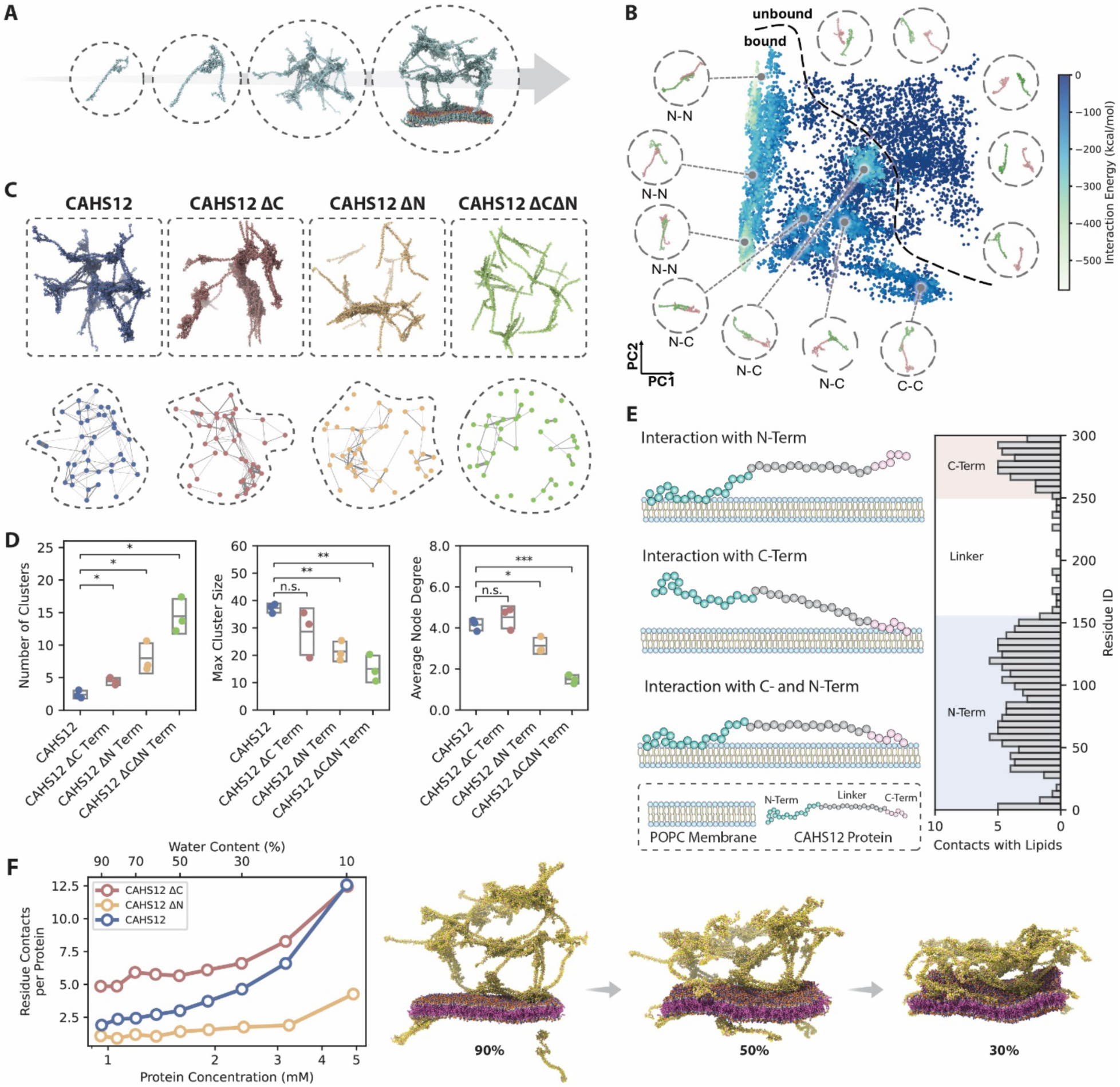
Molecular dynamics simulations of CAHS12 protein assemblies and their interaction with phospholipid membranes. (**A**) Schematic depiction of the simulation systems with increasing structural complexity, ranging from a single CAHS12 monomer (left) to oligomers, fibrillar networks, and large-scale assemblies interacting with a POPC lipid membrane (right). (**B**) A PCA plot of CAHS12 dimer conformations from unbiased simulations presenting representative dimer configurations from selected clusters. The color scale indicates interaction strength based on short-range inter-protein interaction energy. The dashed line approximately delineates the bound and unbound states. (**C**) Network representations of protein assemblies for wild-type CAHS12, and the ΔC, ΔN, and ΔCΔN variants. Nodes represent individual proteins; edges represent the contact among proteins; edge thickness is proportional to the inverse of the center-of-mass distance between two proteins. (**D**) Quantification of differences in topological organization between the CAHS12, CAHS12 ΔC, CAHS12 ΔN, and CAHS12 ΔCΔN variants for the systems shown in (C). The number of clusters represents the count of distinct and connected protein assemblies. The maximum cluster size denotes the largest number of proteins within a single connected cluster. The average node degree indicates the mean connectivity per protein. The box bounds represent standard deviations from three independent replicates, and the middle lines represent the mean. Statistical significance was assessed using Welch’s t-test. (**E**) Schematic models of membrane-binding modes for the CAHS12 protein, illustrating the interaction between different domains of the protein, particularly the C- and N-terminal domains, with the POPC membrane (left). Contact frequency of residues across the whole CAHS12 sequence supporting the proposed models (right). Contacts are quantified by identifying residues within 0.5 nm of POPC lipid headgroups and averaging the counts over simulation frames. (**F**) Residue-level membrane contact analysis for CAHS12, CAHS12 ΔC, and CAHS12 ΔN during progressive water removal simulations (i.e., increasing protein concentration), along with the molecular visualization showing the CAHS12 protein and POPC membrane, mimicking the desiccation process. The percentage indicates the fraction of remaining water beads relative to the initial simulation box, with a lower value corresponding to a greater extent of water removal. Contacts are defined as the number of unique residues that are within 0.5 nm of POPC lipid headgroups, normalized by the number of CAHS12 proteins. All data in (C-F) were collected from the final 500 ns of the 1500 ns simulations. Statistical analyses were performed on averaged data points from three independent simulations. A Welch’s t-test (two-tailed) between CASH12 and CAHS12 ΔC or CAHS12 ΔN was used in (D) to assess statistical significance. *, *p* < 0.05; **, *p* < 0.01.

To further dissect the roles of each terminal region in CAHS12 assemblies, we constructed ΔC and ΔN variants by truncating the respective regions while preserving the conformation of the remaining protein (**Supplementary Fig. 15, Supplementary Table 1, 4, 5, 6**). Network analyses of multimer simulations with POPC membranes shows that both the wild-type CAHS12 and the ΔC variant form protein networks and filaments (**Fig. 2C** and **Supplementary Fig. 16**) with comparable cluster numbers, sizes, and node connectivity (**Fig. 2D**). In contrast, the ΔN variant exhibits a significantly reduced capacity to form networks (**Fig. 2C** and **Supplementary Fig. 16**), with increased cluster numbers and decreased cluster sizes and connectivity (**Fig. 2D**). The ΔCΔN variant leaving just the linker region leads to a further decrease in capacity to form interconnected assemblies. These results highlight the critical role of the N-terminal region as a “sticky” end that promotes aggregation and filament formation, potentially serving as mechanical support for membrane architectures during dehydration, and providing a putative rationale for the experimentally observed role for the N-terminus in conferring protective function (**Fig. 1B**, **Supplementary Fig. 6**). These simulation results agree with prior experimental observations from prior studies demonstrating that CAHS protein mutants can form diverse molecular assemblies and confer protective effects^9,10^. In addition to mediating protein-protein interactions, we hypothesized that the CAHS12 protein may also mediate protein-lipid interactions, anchoring or stabilizing membrane structures under desiccation. Such an idea has been proposed for tardigrade-disordered proteins by Bino *et al.* and Hesgrove *et al.*^7,29^. To test this, we analyzed contact frequencies between different regions of CAHS12 and the lipid membrane using simulations of the protein network in the presence of a POPC bilayer. Residues, such as serine, threonine, and glycine, in both the N- and C-terminal regions are frequently localized to or embedded within the lipid surface, whereas the linker region showed markedly fewer membrane contacts (**Fig. 2E and Supplementary Fig. 17A**). Trajectory analysis revealed three distinct protein–membrane interaction modes, involving membrane association mediated by the N- and C-terminal regions and the proximity of the α-helix linker to the membrane (**Supplementary Fig. 18**), suggesting a variety of mechanisms by which CAHS12 engages with lipid structures. Prior studies have shown that the interaction patterns between CAHS12 proteins and phospholipid membranes are sensitive to membrane composition^30,31^. Increasing the fraction of the anionic lipid POPG in the membrane enhances the contact frequency of the linker region while preserving the high contact frequency of the terminal regions (**Supplementary Fig. 19**), indicating that the terminal domains, along with the linker domain, cooperate in the interaction with lipid membrane^32^. In this study, we focus on POPC membranes in that they offer a well-defined reference system for investigating the utility of CAHS12 in the context of synthetic cells with POPC membranes. As such, our simulation results are intended to characterize CAHS12 interactions with neutral POPC membranes and are not necessarily transferable to the understanding of CAHS12 interactions with natural biological membranes.

We next simulated dehydration by sequential deletion of water beads in simulations involving CAHS12, CAHS12 ΔN, and CAHS12 ΔC variants in the presence of a POPC membrane. Interestingly, while the CAHS12 ΔC variant exhibited 2-fold higher membrane contacts in the hydrated state (**Supplementary Fig. 17B**), contact frequencies of the CAHS12 ΔC variant and full-length CAHS12 increase to a comparable level under dehydrated conditions (**Supplementary Fig. 20)**. In contrast, the CAHS12 ΔN variant displays a markedly reduced ability to interact with the membrane, further emphasizing the critical role of the N-terminal region in membrane association (**Fig. 2F**, **Supplementary Fig. 20**). These simulation results are in good alignment with experimental findings that both full-length CAHS12 and the ΔC variant protect GUV morphology during dehydration, whereas the ΔN variant fails to do so (**Fig. 1B**, **Supplementary Fig. 6**). Visualization of simulation trajectories further confirms that CAHS12 aggregates and attaches to the membrane surface, which is consistent with experimental observations of a proteinaceous layer forming on membranes during dehydration (**Fig. 1G**). Collectively, these results elucidate a dual role for the CAHS12 protein, particularly its N-terminal region, in driving both protein network formation and membrane association, thereby providing mechanistic insight into how CAHS12 may contribute to membrane stabilization during desiccation.

### CAHS12 preserves GUV integrity during long-term storage and under environmental stress

To assess whether CAHS12 can confer long-term membrane stability, we monitored GUV morphology following ambient storage under both hydrated and desiccated conditions. GUVs encapsulating CAHS12-GFP retained their size and spherical shape after 5 to 14 days at room temperature (p > 0.05, n > 100). Even after 30 and 74 days, vesicles displayed only a slight size reduction but maintained distinct spherical geometry (**Fig. 3A, Supplementary Fig. 21**). In contrast, GUVs lacking CAHS12 or containing the ΔN variant deformed or disappeared after just 5 days of storage, accompanied by cargo leakage; by 14 days, intact vesicles were no longer detectable or sparse (**Fig. 3A, Supplementary Fig. 21**). These findings indicate that CAHS12 substantially delays the structural deterioration of GUVs, likely through interfacial interactions that buffer mechanical stress from water loss or chemical aging.

**Fig. 3.**
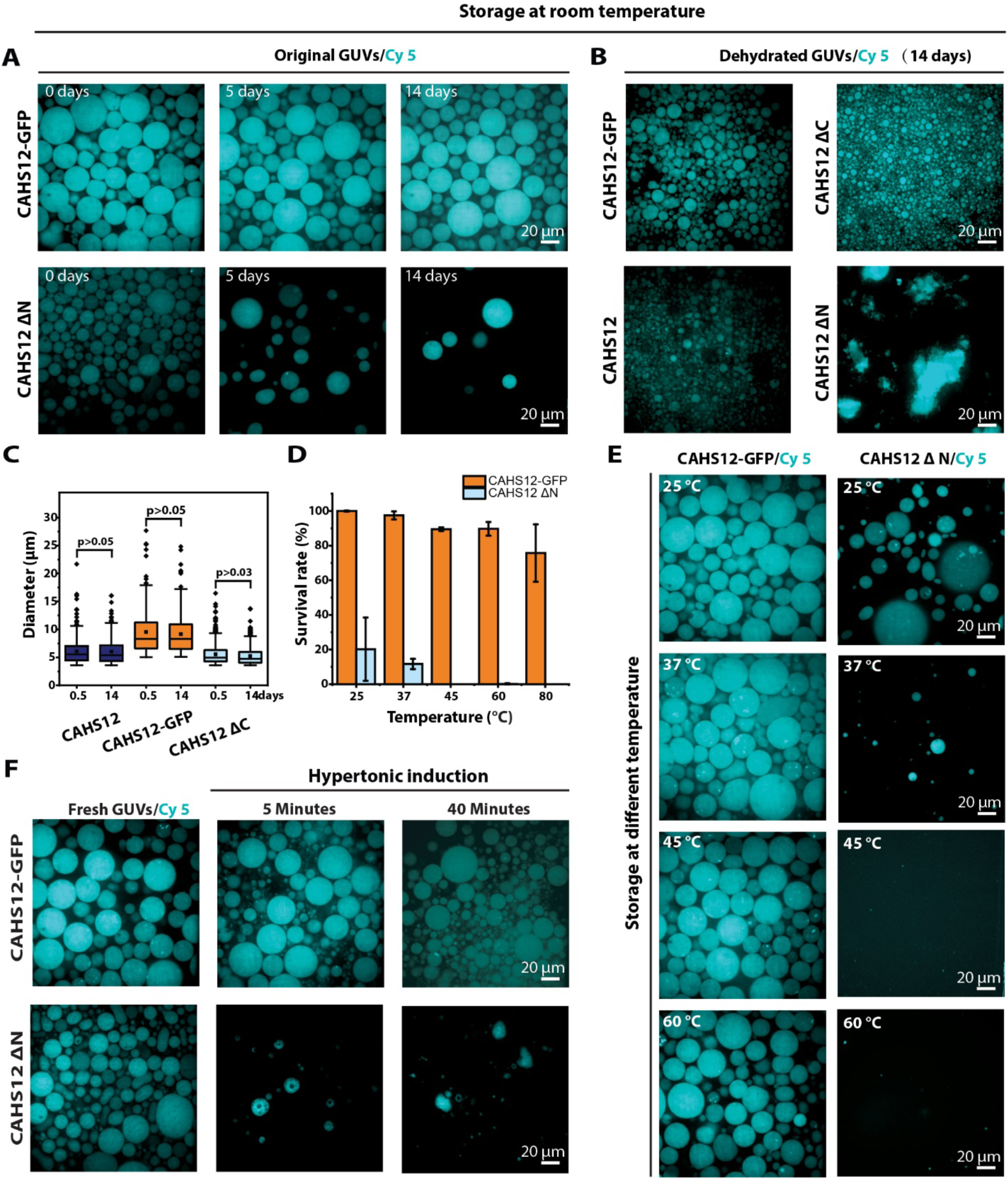
CAHS12 enhances GUV stability during storage, dehydration, and osmotic stress. (**A**) Confocal images of GUVs containing Cy5 stored at room temperature. GUVs containing 280 µM CAHS12-GFP or CAHS12 ΔN were stored for 0–14 days. (**B**) Confocal images of GUVs containing 280 µM CAHS12 variant after continuous dehydration and storage for 14 days. (**C**) Diameter distributions of GUVs containing CAHS12, CAHS12-GFP, or ΔC over continuous dehydration for 0.5 and 14 days. Retention rates (**D**) and confocal images (**E**) of GUVs after incubation at increasing temperatures (25–60 °C), where 25 °C was stored for 5 days, 37 °C and 45 °C was stored for 2 days, 60 °C was stored for 2 hours. (**F**) Fluorescence microscopy images of GUVs in an equal volume high-osmolarity glucose solution (1200 mOsm/kg) at the indicated times, where the external solution has an osmotic pressure 2.5 times that of the GUV interior.

We next examined whether CAHS12 could stabilize GUVs under prolonged desiccation, mimicking long-term dry storage. GUVs encapsulating full-length CAHS12, CAHS12-GFP, or ΔC variants retained spherical morphology and high vesicle density after 14 days of dehydration at room temperature, with minimal aggregation or rupture (**Fig. 3B**). Notably, vesicle size did not change significantly between 0.5 and 14 days of desiccation (p > 0.05), suggesting that CAHS12 maintains membrane architecture even under continuous drying (**Fig. 3C**). In contrast, GUVs containing ΔN variant showed pronounced collapse and membrane failure. To evaluate the robustness under physical stress, we subjected GUVs to heat and osmotic shock—common environmental stressors that challenge membrane integrity. When exposed to temperature from 25 to 80 °C under ambient humidity, GUVs containing CAHS12-GFP retained spherical morphology and high vesicle abundance across the full range, with retention rate exceeding 80% (**Fig. 3D-E**). In contrast, ΔN-containing or empty GUVs exhibited structural failure even at moderate temperatures (37 °C), with near-complete loss above 45 °C (**Supplementary Fig. 22**). These results underscore the essential role of the N-terminal region in thermal protection and demonstrate an unusually broad thermal operating window relative to traditional protein-based stabilizers. To test CAHS12’s ability to buffer against acute osmotic stress (Infusion by an equal volume of a 1200 mOsm/kg glucose solution), we exposed GUVs to a high-osmolarity solution and imaged their morphology at 5- and 40-minutes post-treatment (**Fig. 3F**). CAHS12-GFP–containing GUVs resisted collapse throughout the exposure, maintaining circular geometry. By contrast, CAHS12 ΔN–containing vesicles rapidly lost structure, showing shrinkage, deformation, and rupture within 5 minutes (**Supplementary Fig. 23**). At 40 minutes, few intact vesicles remained. These findings again support a model in which CAHS12 assembles into a dynamic, stress-responsive protein coating that preserves membrane shape and elasticity under compression. This membrane-centric mechanism may underlie the exceptional survival capacity of tardigrade cells under desiccation and extreme conditions. Moreover, the ability to preserve vesicle architecture under ambient and dry storage conditions provides a foundation for engineering cold-chain–independent synthetic cells and bioformulations with extended shelf life.

### Robust preservation and rehydration-triggered activation of cell-free systems

When a tardigrade experiences water scarcity, it enters a dormant state called a tun, losing up to 97% of its body water^33^. Encouraged by CAHS12’s ability to protect synthetic membranes against a range of environmental stresses, we sought to determine if synthetic cells with CAHS proteins could enter a tun state under desiccation. Synthetic cell construction represents a frontier of biological engineering, with the eventual goal of building a functioning cell ‘from scratch’^34–36^. Our synthetic cells consist of GUVs encapsulating cell-free transcription–translation (TX-TL) machinery based on the PURE (Protein synthesis Using Recombinant Elements) system^37^ and DNA encoding genes of interest. As illustrated in **Fig. 4A**, we hypothesize that CAHS proteins protect synthetic cells against dehydration to arrest biochemical reactions while retaining membrane structure, and rehydration restores cellular functions. Dehydration halts cell-free expression (CFE) activity by water removal, while CAHS proteins preserve the internal contents in a dormant yet structurally intact configuration. Upon rehydration, the previously inactivated TX-TL processes would be reactivated, initiating *de novo* CFE within the synthetic cells. Notably, encapsulation within synthetic cells provides an additional layer of protection during desiccation by physically separating the reaction components from external degradation factors such as proteases and RNases, an advantage not afforded by dehydrating reactions in bulk solution or on paper. Upon rehydration, water re-entry dissolves the CAHS networks, restores molecular mobility, reactivates the previously halted TX-TL machinery, and initiates *de novo* CFE within the synthetic cells. This enables synthetic cells to toggle between dormant and active states while maintaining functional integrity under extreme water loss and environmental stress.

**Fig. 4.**
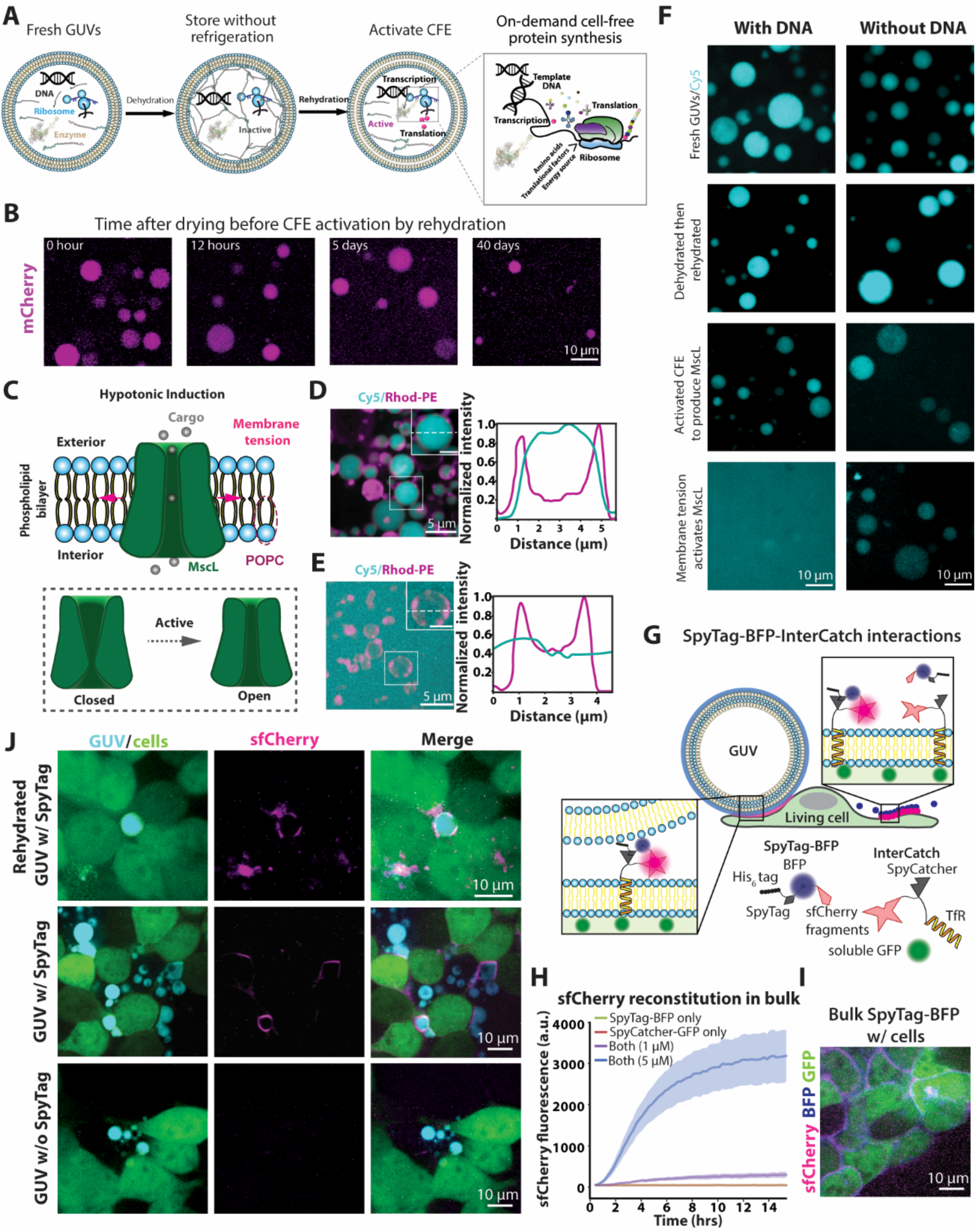
Long-term preservation and activation of CFE and membrane protein function in CAHS12-stabilized GUVs. (**A**) Schematic of the workflow: synthetic components, including DNA and TX-TL machinery, are encapsulated in GUVs, dehydrated for dry storage, and rehydrated to trigger cell-free expression. (**B**) Fluorescence microscopy images of CFE activated by rehydration of dehydrated GUVs stored at room temperature for different periods (0 to 40 days). (**C**). Schematic of MscL activation via hypotonic induction, which creates elevated membrane tension and promotes channel opening. (**D–E**) Fluorescence microscopy images and corresponding line-scan intensity profiles showing localization of Cy5 cargo and Rhod-PE membrane dye before (**D**) and after (**E**) MscL activation. Redistribution of Cy5 signal indicates membrane permeability upon functional channel activation. (**F**) Functional demonstration of MscL activity: Cy5-containing GUVs with or without DNA encoding MscL were subjected to drying, rehydration, CFE activation, and hypo-osmotic shock. Only GUVs with MscL DNA show dye leakage upon osmotic induction. **(G)** Schematic illustrating SpyTag-BFP and InterCatch-mediated interactions between synthetic cells (GUVs) and living cells, resulting in sfCherry reconstitution. SpyTag-BFP, SpyTag attached to Tag-BFP and one fragment of split sfCherry, is displayed on the GUV membrane. InterCatch, consisting of SpyCatcher fused to the complementary sfCherry fragment and the transmembrane domain of transferrin receptor (TfR), is stably expressed on HEK293T cell membranes. Soluble GFP is co-expressed for visualization. **(H)** *In vitro* sfCherry reconstitution in bulk solution using purified SpyTag-BFP and SpyCatcher-GFP. **(I)** sfCherry fluorescence is observed when InterCatch cells are incubated with purified SpyTag-BFP. **(J)** Following incubation, sfCherry reconstitution between GUV and cells is observed with both freshly prepared and dehydrated/rehydrated SpyTag-BFP GUVs. No reconstitution is seen with GUVs lacking SpyTag-BFP.

We first assessed whether CAHS12 can preserve enzymatic function through dehydration–rehydration cycles, by encapsulating horseradish peroxidase (HRP) in GUVs and subjecting the system to drying for 12 hours. Upon rehydration, HRP was activated by the addition of hydrogen peroxide, initiating a catalytic reaction. The appearance of the fluorescent product resorufin within the GUVs confirmed that enzymatic activity was retained after desiccation (**Supplementary Fig. 24**), indicating that CAHS12 enables functional preservation of encapsulated proteins under extreme drying stress. Prior studies have established that CAHS proteins can act as protein stabilizers during drying, preserving enzymatic activity *in vitro*^9,32,38,39^. Our results confirm and extend these findings by demonstrating that sequence-based variants of CAHS differ in their efficacy across client molecules and environmental conditions, suggesting mechanisms by which intrinsic disorder and phase behavior tune desiccation protection.

It has been previously demonstrated that CFE reactions can be freeze-dried onto paper or in a tube to enable diagnostic applications or on-demand biomanufacturing by simply adding water and DNA^40,41^. However, synthetic cells are not expected to maintain structural and functional integrity upon dehydration without protection, similar to our test in *E. coli* (**Supplementary Fig. 1**). To explore whether CAHS proteins can protect synthetic cells from desiccation stress, we encapsulated CFE components and mCherry-expressing DNA (as a fluorescent reporter) and then evaluated how dry storage duration affects CFE output (**Fig. 4B**). Synthetic cells containing CAHS12-GFP were dehydrated and stored at room temperature for defined time intervals, then rehydrated and incubated at 37 °C to initiate protein expression. After dehydration and rehydration immediately (0 h) to activate TX-TL, we observed strong mCherry fluorescence across nearly all vesicles, confirming efficient preservation of transcriptional and translational activity. After 12 hours of dry storage, synthetic cells maintained structural integrity and expressed mCherry at comparable levels. Even after 5 days of storage, rehydrated GUVs maintained high expression levels, demonstrating sustained biochemical activity. Remarkably, after 40 days or even 60 days of ambient storage, GUVs still preserved morphology and measurable CFE activity, though expression levels became heterogeneous. In contrast, and as expected, synthetic cells lacking CAHS12-GFP did not activate CFE upon rehydration (**Supplementary Fig. 25**). These findings indicate a gradual decline in system performance over time but highlight the surprising durability of both lipid membranes and encapsulated TX-TL machinery under non-refrigerated, dry storage conditions.

The successful rebooting of CFE in synthetic cells following prolonged desiccation prompted us to ask whether other synthetic cell functions can be preserved. We are particularly interested in membrane-active properties, as most cell-cell communication takes place at the membrane interface via membrane proteins. As a proof-of-concept demonstration, we synthesized the bacterial mechanosensitive channel^42–44^, MscL, *de novo* via CFE inside synthetic cells under different conditions, which can be subsequently activated by membrane tension to mediate cargo release. The working model (**Fig. 4C**) depicts MscL as a pentameric transmembrane pore that responds to hypotonic stress by sensing membrane tension^45^. Upon rehydration and followed by hypo-osmotic induction, water influx generates lateral tension across the synthetic cell membrane, triggering MscL opening and rapid efflux of encapsulated cargo. To test this, desiccation-resistant synthetic cells co-encapsulating a fluorescent cargo (i.e., Cy5) and membrane label (i.e., Rhod-PE) underwent CFE expressing MscL, followed by dehydration–rehydration to test MscL functionality. In hypertonic conditions, Cy5 remained well-encapsulated, as expected (**Fig. 4D** and **Supplementary Fig. 26**). Under hypotonic stress, Cy5 fluorescence rapidly dissipated from the vesicle interior (**Fig. 4E** and **Supplementary Fig. 26**), with increased background signal and flattening of radial gradients, indicating leakage. Rhod-PE signal confirmed that the membrane was preserved while permeability was altered. These results support successful membrane integration and mechanical gating of MscL to trigger cargo release.

To dissect the sequence of events required for successful MscL-mediated release, we compared desiccation-resistant synthetic cells with or without MscL-encoding DNA after CFE, dehydration–rehydration, and hypotonic challenge (**Fig. 4F**). In freshly prepared synthetic cells, both conditions retained Cy5 cargo. After dehydration for 12 hours and rehydration, synthetic cell morphology and content remained intact. Rehydration in the presence of CFE reagents resulted in pronounced Cy5 loss following hypotonic exposure only in synthetic cells containing MscL DNA, confirming that membrane tension alone was insufficient to induce leakage in the absence of MscL. These findings demonstrate that synthetic cells can be used as long-lived, rehydration-activated chassis for synthetic gene expression and functional protein integration.

To determine whether CAHS12 preserves membrane protein function following dehydration and rehydration, we tested a model of membrane-mediated communication between synthetic and living cells using a juxtacrine signaling system based on SpyTag/SpyCatcher conjugation^46^. In this platform, HEK293T cells stably express a membrane-anchored SpyCatcher fused to one half of split superfolder Cherry (sfCherry1–10), hereafter referred to as InterCatch, served as the living cell component. Synthetic cells were functionalized with SpyTag fused to a blue fluorescent protein and the C-terminal sfCherry fragment (sfCherry11) to form a SpyTag–BFP–sfCherry11 fusion. This fusion was anchored to the synthetic membrane via Ni²⁺–His₆ coordination to surface-displayed lipids containing nickel–nitrilotriacetic acid (Ni–NTA).

As a validation step, we first confirmed successful sfCherry complementation in solution using purified SpyTag–BFP–sfCherry11 and InterCatch proteins, which yielded robust red fluorescence upon coincubation, indicating functional interaction between the split fragments (**Fig. 4H** and **Supplementary Fig. 27A-B**). sfCherry complementation is also preserved in HEK293T cells expressing InterCatch-GFP incubated with soluble SpyTag-BFP (**Fig. 4I**). We then tested whether synthetic cells could support this interaction following stress. SpyTag–BFP–functionalized synthetic cells (contained 100 μM CAHS12) dehydrated under vacuum conditions (12 h at 25 °C), and subsequently rehydrated in PBS. Rehydrated synthetic cells were applied to a monolayer of adherent HEK293T–InterCatch cells and incubated at 37 °C for 4 hours. Fluorescence microscopy revealed red sfCherry signal localized at the interface between synthetic and living cells, consistent with membrane-proximal complementation of the split protein (**Fig. 4J** and **Supplementary Fig. 27C**). Control experiments confirmed that the interaction was specific and dependent on both the membrane display of SpyTag and the presence of CAHS12 (**Fig. 4J**). No sfCherry fluorescence was observed when synthetic cells lacking SpyTag–BFP were applied to InterCatch cells, regardless of dehydration state. Similarly, dehydrated synthetic cells lacking CAHS12 showed little to no synthetic cell formation upon rehydration (**Supplementary Fig. 28)**, and no sfCherry signal was observed after incubation with InterCatch cells, underscoring CAHS’s ability to preserve membrane protein display and functional architecture under desiccation stress. Our results reveal that the tardigrade-derived protein CAHS12 acts as a membrane-active, intrinsically disordered polymer that preserves synthetic cell architecture under extreme physical stress. Through its N-terminal region, CAHS12 engages lipid interfaces, forming a dynamic, reversible coating that buffers dehydration-induced membrane stress, prevents rupture, and enables rapid structural recovery upon rehydration. Structural, biophysical, and computational analyses converge on a mechanism wherein CAHS12 stabilizes vesicle shape and integrity by modulating interfacial energetics and assembling into stress-responsive supramolecular networks.

The global biotechnology sector remains tethered to cold-chain logistics^47–49^. From mRNA vaccines to engineered microbes, nearly all bioactive materials require refrigeration or freezing to maintain efficacy during storage and transp^50^. This dependence limits deployment in remote or resource-constrained regions. The protective function by CAHS proteins extends to cell-free gene expression systems: dehydrated CAHS12-stabilized synthetic cells retain biosynthetic capacity after long-term ambient storage and are reactivated by simple rehydration. Reconstituted mechanosensitive channels further enable stimulus-triggered molecular release, mimicking biological secretory functions. These findings establish a modular platform for engineering desiccation-tolerant synthetic cells with programmable behaviors, opening new avenues for field-deployable biosystems, cold-chain–independent biomanufacturing, and adaptive biointerfaces.

## Methods

### Protein expression and purification

Full-length CAHS12 from *Ramazzottius varieornatus* (a gift from Takekazu Kunieda, University of Tokyo) and truncation variants (CAHS12-GFP, -BFP, ΔN, and ΔC) were cloned into the pET-28b(+) vector (Novagen) with an N-terminal His_6_-tag for affinity purification. Constructs were verified by sequencing and transformed into Escherichia coli BL21(DE3) cells (New England Biolabs)^51^. A single colony was inoculated into 5 mL of LB broth supplemented with 50 μg mL⁻¹ kanamycin and cultured overnight at 37°C with shaking at 220 rpm (Thermo Scientific, Solaris shaker). The overnight culture was diluted 1:200 into 1 L of fresh LB medium with kanamycin and grown at 37°C until the optical density at 600 nm (OD₆₀₀) reached 0.5–0.6. Cultures were rapidly cooled to room temperature in an ice-water bath and induced with 0.2 mM isopropyl β-D-1-thiogalactopyranoside (IPTG, Thermo Fisher). Expression proceeded at 24°C for 16 h with shaking.

Cells were harvested by centrifugation at 6,000 × g for 30 min at 4°C and resuspended in lysis buffer (20 mM Tris-HCl, pH 8.0, 500 mM NaCl, 5 mM imidazole). The suspension was lysed by sonication on ice and clarified by centrifugation at 25,000 × g for 30 min at 4°C. Supernatants were filtered (0.45 µm) and loaded onto a 1 mL HisTrap FF column (Cytiva) using an ÄKTA Start FPLC system (Cytiva). The column was washed with 20 column volumes of wash buffer (20 mM Tris pH 8.0, 500 mM NaCl, and 25 mM imidazole) and eluted with a solution of 250 mM imidazole in 20 mM Tris-HCl (pH 8.0), 500 mM NaCl. Fractions (1.5 mL each) were collected at a flow rate of 1 mL min⁻¹, and protein purity was assessed by SDS-PAGE. Peak fractions were pooled and dialyzed overnight at 4°C against 20 mM Tris-HCl (pH 8.0), 25 mM NaCl, and 1 mM EDTA using a 10 kDa molecular weight cutoff membrane (Spectrum Labs). Dialyzed protein was aliquoted, flash-frozen in liquid nitrogen, and stored at –80°C until use. Amino acid sequences of each construct are provided in **Supplementary Tables 1–5**.

### Giant unilamellar vesicle (GUV) preparation by phase transfer

GUVs were prepared using a modified phase transfer protocol optimized for high encapsulation efficiency and membrane uniformity^52,53^. Lipid films were generated by dissolving a lipid mixture of 99.9 mol% 1-palmitoyl-2-oleoyl-sn-glycero-3-phosphocholine (POPC, Avanti) and 0.1 mol% Rhodamine B-labeled phosphoethanolamine (Rhod-PE) in chloroform to a final concentration of 0.41 mM. The lipid solution was deposited in a clean glass vial and evaporated under a gentle argon stream while rotating the vial to ensure uniform film distribution. Residual solvent was removed by placing the vial under vacuum for ≥1 hour. The dried lipid film was resuspended in 1.1 mL of mineral oil (Sigma) by vortexing for 30 seconds and incubated at 55–75 °C for 20 minutes in a dry chemical oven, with vortexing every 10 minutes to promote full lipid dispersion. To form a stable oil–water interface, 400 µL of outer aqueous solution (300 mM glucose) was added to a 1.5 mL microcentrifuge tube, and 300 µL of the lipid–oil solution was carefully layered on top using a slow pipetting motion against the inner wall to minimize turbulence. The interface was allowed to equilibrate for 1 hour at room temperature. Separately, 20 µL of inner aqueous solution (300 mM sucrose), containing proteins, Cy5 (10 µg/mL), was added to a fresh tube. This was gently overlaid with 600 µL of the lipid–oil mixture, and the two phases were emulsified by slow pipetting (≥100 strokes or until the emulsion became uniformly colored). The emulsion was then slowly added onto the preformed oil–water interface. The assembled gradient was centrifuged at 2500 × g for 10 min at room temperature using a swing-bucket rotor (Eppendorf, 5415R). Following centrifugation, the upper oil phase and aggregated material were carefully aspirated using a 200 µL pipette. Once the remaining oil volume reached ∼100 µL, finer removal was performed using a 10 µL pipette tip in a slow circular motion, changing tips to avoid contamination. The GUV pellet at the bottom of the tube was resuspended by gentle pipetting (10–15 times) until the solution appeared homogeneously colored. GUVs were immediately transferred to optical polymer bottomed imaging plates (MatTek) by pipetting against the well bottom in a circular pattern. GUV integrity and encapsulation efficiency were confirmed by confocal fluorescence microscopy prior to downstream analyses.

### Dehydration–rehydration cycling

5 µl solution of GUVs was transferred into 96-well glass-bottom plates. To initiate desiccation, plates were placed in a vacuum desiccator and incubated under ambient temperature (22–25°C) at ∼31 mbar for ≥12 h to ensure complete water removal. Rehydration was carried out by gently adding 5 μL of ultrapure deionized water to each well without disturbing the dried film. Samples were incubated at room temperature for 10 min to allow for complete rehydration. Vesicle morphology, integrity, and size distribution were analyzed by confocal fluorescence microscopy (Yokogawa CSU-X1) using Rhodamine-PE and/or encapsulated fluorescent cargo as reporters. Z-stacks were acquired for 3D reconstruction where indicated.

### Interfacial tension measurements

Interfacial tension (IFT) between aqueous protein solutions and oil phases was quantified using a pendant drop tensiometer (Krüss DSA100)^16^. The continuous phase consisted of mineral oil with or without 0.41 mM 1-palmitoyl-2-oleoyl-sn-glycero-3-phosphocholine (POPC) to assess lipid-specific interfacial interactions. A stainless-steel needle (outer diameter: 0.904 mm) was vertically positioned in the measurement cuvette and filled with 5 μL of aqueous solution containing CAHS12 or variant proteins at defined concentrations (0–1 mM). Care was taken to eliminate all air bubbles from the syringe and needle prior to droplet formation. Once the droplet was formed, it was allowed to equilibrate in the continuous phase for 30 min to ensure full adsorption and interfacial stabilization. Image acquisition was performed in real time, and interfacial tension values were calculated using the Young–Laplace equation via the instrument’s built-in analysis software. For each condition, IFT values were averaged from three independent measurements. Only droplets displaying stable geometry and no drift during the equilibration period were included in the analysis.

### Cell-free expression and encapsulation in synthetic vesicles

Cell-free protein synthesis was performed using a reconstituted *in vitro* coupled transcription/translation system (PUREfrex 2.0, GeneFrontier Corporation) with minor modifications to enable encapsulation within GUVs and compatibility with CAHS12-containing systems. All assembly steps were conducted on ice to minimize enzymatic degradation and preserve reaction activity. Each 20 µL reaction mixture was prepared in low-retention microcentrifuge tubes (Axygen) and contained: 10 µL of Solution 1 (providing Amino acids, NTPs, tRNAs and substrates for enzymes etc.), 1 µL of Solution 2 (Proteins in 30% glycerol buffer), 2 µL of Solution 3 (ribosomes, 20 µM), and 1 µL of RNase inhibitor (40 U/µL, New England Biolabs) to protect mRNA during the reaction. To facilitate the subsequent formation of GUVs, add 1.2 µL of OptiPrep (Sigma-Aldrich) to form a gentle density gradient so that the droplets containing the above solution can penetrate the oil-water interface under centrifugal force.

For mechanostability and stress-response testing, recombinant CAHS12 protein (3.8 µL, final concentration: 0.19 mM) was added to each reaction immediately before the DNA template. Plasmid DNA encoding either soluble mCherry or the mechanosensitive channel MscL (homemade, purified using Qiagen miniprep kits) was added in amounts of 2 ng/µL per 1 kb per reaction, with precise quantities optimized for each plasmid. Ultrapure nuclease-free water (Invitrogen) was added last to adjust the total volume to 20 µL. For bulk CFE measurements, reactions were transferred into a black-walled, clear-bottom 96-well plate. Plates were sealed with optically clear adhesive film to prevent evaporation. Fluorescence (Ex/Em: 580/610 nm for mCherry) was monitored every 2 minutes at 37 °C over 8 hours using a BioTek Synergy H1 plate reader.

To prepare GUVs encapsulating CFE systems, the phase transfer method was employed with the following modification: the inner aqueous phase consisted of the complete 20 µL CFE reaction described above (osmolarity ∼1079 mOsm/kg), while the outer solution comprised Tris-HEPES buffer (pH 7.4), adjusted to ∼1040 mOsm/kg. To prevent premature initiation of protein synthesis or osmotic imbalance, all encapsulation steps were performed at 4 °C or on ice. Following centrifugation (2500 × g, 10 min, 4 °C), vesicles were collected and stored on ice. Protein expression within GUVs was initiated by warming to 37 °C.

Vesicle fluorescence was imaged using confocal microscopy (Yokogawa CSU-X1) at defined time points. GUVs displaying uniform internal fluorescence were classified as functionally expressing synthetic cells.

To assess the functional activity of reconstituted MscL within GUVs, osmotic downshock was applied to induce membrane tension and trigger channel opening. Following CFE incubation, GUVs containing MscL and an encapsulated fluorescent dye (e.g., Cy5, 10 µM) were transferred to imaging chambers (96-well plate). Vesicles were initially imaged in isotonic buffer (e.g., 1079 mOsm/kg CFE solution/Tri-HEPES solution) to record baseline fluorescence. For activation, the external buffer was rapidly exchanged with a hypo-osmotic solution (∼500 mOsm/kg, prepared by diluting the isotonic buffer with Milli-Q water) while continuously imaging using confocal microscopy to monitor dye efflux.

Fluorescence loss was quantified by measuring the mean internal fluorescence intensity within individual GUVs over time using Fiji (ImageJ), normalized to the initial intensity prior to downshock. MscL-mediated permeability was confirmed by the rapid decrease in fluorescence upon hypo-osmotic challenge compared to control vesicles lacking MscL. Multiple vesicles (>30 per condition) were analyzed across at least three independent preparations to confirm reproducibility of MscL activation and functional pore formation under mechanical stress.

### GUV–living cell functional interaction via SpyTag–SpyCatcher binding

HEK293T cells stably expressing membrane-bound SpyCatcher and soluble GFP were previously developed^51^. Cells were cultured in DMEM (Gibco) supplemented with 10% fetal bovine serum (Gibco), 1% penicillin–streptomycin (Gibco), and maintained at 37 °C in a humidified incubator with 5% CO₂. Puromycin (2 μg/mL; Gibco) was added to ensure robust expression of SpyCatcher and GFP.

SpyTag-BFP (SpyTag–BFP–sfCherry11) and SpyCatcher-GFP (SpyCatcher–GFP–sfCherry1-10) proteins were purified according to previously published protocols^51^. For bulk sfCherry reconstitution assays, 10 μL reactions containing SpyTag-BFP (5 μM), SpyCatcher-GFP (5 μM), or both proteins (1 μM or 5 μM each) were prepared in glass-bottom 384-well plates (Corning). Fluorescence was monitored using a Cytation 5 imaging plate reader (BioTek) at room temperature every 15 minutes for 15 hours (Ex/Em: 580/615 nm).

GUVs were prepared using the inverted emulsion method with the following parameters: (1) The membrane composition consists of 92% POPC, 5% 18:1 1,2-dioleoyl-sn-glycero-3-[(N-(5-amino-1-carboxypentyl)iminodiacetic acid)succinyl] (nickel salt) (DGS-NTA(Ni)), and 3% cholesterol conjugated to polyethylene glycol 2000 (Chol–PEG2k). (2) The GUV inner solution was prepared using a 330 mOsm/kg solution of sucrose and contained 6% OptiPrep, 10 μM Cy5, and 100 μM His-tag cleaved CAHS12. (3) The outer solution consists of phosphate-buffered saline (PBS) (∼300 mOsm/kg).

After fabrication, SpyTag-BFP (2 μM) was added to the GUV solution and incubated for 15 minutes before addition to cells or dehydration via desiccation. For bulk interactions between SpyTag-BFP GUVs and soluble SpyCatcher-GFP, 2 μM SpyCatcher-GFP was added to GUVs and imaged both immediately and after 4 hours at room temperature. For interactions between SpyCatcher-expressing cells and soluble SpyTag, 1 μM SpyTag was added to cells cultured on glass-bottom dishes and imaged immediately and after 4 hours.

For GUV–cell interaction experiments, SpyCatcher-expressing HEK293T cells (5×10^5^) were seeded in 6-well plates with a 20 mm glass-bottomed micro-well (Cellvis) and allowed to adhere for 24 hours. Before the experiment, the media was aspirated and replaced with 100 μL PBS. Then, 50 μL of SpyTag-labeled GUVs was added, and the plate was centrifuged at 300 × g for 15 minutes to promote GUV-cell contact. Cells were incubated with GUVs at 37 °C for 4 hours prior to imaging.

All imaging was conducted on a spinning-disk confocal microscope, using the same setup described previously, except with the following exposure times: BFP (405 nm, 500 ms), GFP (488 nm, 1,000 ms), sfCherry (561 nm, 2,000 ms), and Cy5 (640 nm, 100 ms). Images were processed using Fiji (ImageJ), with brightness and contrast adjusted for visualization.

### Molecular dynamics simulations

#### Computational Modeling of a CAHS12 Protein

To investigate the interaction between the CAHS12 protein and a POPC membrane, we conducted coarse-grained (CG) molecular dynamics (MD) simulations using the Martini3 force field^54^. While all-atom simulations provide higher resolution and accuracy, their computational cost limits their applicability for probing large-scale protein self-assembly. In contrast, CG modeling allows simulations of larger systems and time scales, making it suitable for capturing the dynamic assembly behavior of CAHS12. The amino acid sequence of CAHS12 from *Ramazzottius varieornatus* was retrieved from the NCBI database^55^ and converted into an all-atom 3D structure using the AlphaFold3^56^. The resulting structure was subsequently coarse-grained using the martinize2 tool^57^. Given that CAHS12 contains extensive intrinsically disordered regions, appropriate assignment of secondary structure elements during coarse-graining is essential to preserve the protein’s biophysical properties^58^. Previous studies suggest that the CAHS protein adopts a dumbbell-like conformation characterized by disordered N-terminal (residues 1-153) and C-terminal (residues 268-297) regions flanking an ordered α-helical linker domain (residues 154-267) under condensed conditions^10,26^. The local κ values of the sequence – a measure of charge patterning within a protein sequence, with high values favoring disordered conformations^59^ – were computed with a window size of 50 residues using the localCIDER package^60^, which further supports the hypothesized structure (**fig. S10**). The secondary structure predicted by AlphaFold3 was consistent with experimental observations and was used to guide structural assignment via the Define Secondary Structure of Proteins (DSSP) algorithm during model preparation^61,62^. Charges of ionizable residues were assigned under a physiological pH = 7 (E: −1, D: −1, K: +1, R: +1). The final protein model comprises the following secondary structure elements, with C, E, T, S, and H denoting random coils, β-sheets, turns, bends, and α-helices, respectively. For the ΔC and ΔN variants, the respective sequence segments were truncated, while the secondary structure assignments for the remaining regions were preserved.

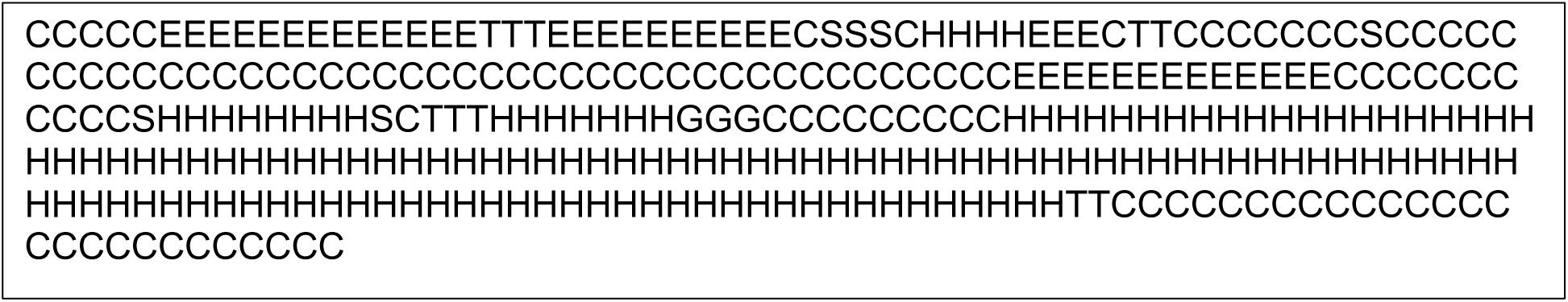

#### Simulations of the CAHS12 Proteins

To determine the radius of gyration (*Rg*) of the CAHS12 protein, a single protein was placed in a cubic water box with a side length of 30 nm, and the net charge was neutralized by the addition of nine sodium ions. The system was first energy-minimized followed by a 40 ns constant pressure and temperature (NPT) equilibration. A subsequent 200 ns production run was performed, and the radius of gyration was computed from the trajectory using the *gmx gyrate* utility in GROMACS. The resulting radius of gyration for a single CAHS12 protein was (5.21 ± 0.11) nm, which is consistent with the range of experimentally determined values reported for other CAHS family proteins^10,27^. To investigate the interaction modes underlying CAHS12 protein assembly into networks, we performed nine independent unbiased simulations with each containing two CAHS12 proteins. The proteins were initially positioned in a 30 nm cubic water box with sufficient separation to prevent direct interactions. Each system underwent energy minimization, followed by 5 ps of constant volume and temperature (NVT) equilibration and 10 ns of NPT equilibration. A 1000 ns production run was then conducted for each replicate to allow for the formation of dimers. To characterize protein-protein interactions, each trajectory frame was featurized using the pairwise distances between α-carbon beads of the two proteins, concatenated with the center-of-mass (COM) distance between them. Principal component analysis (PCA)^63^ over the coordinates of the aligned dimer configurations, followed by k-means clustering^64^ in the top two PCs was employed to identify the landscape of bound dimer conformations. To quantify interaction strength across binding modes, short-range interaction energies between the proteins were computed by rerunning the trajectories with energy decomposition. These analyses revealed that interactions involving the terminal regions predominantly contribute to dimer formation and likely serve as key drivers of protein network assembly.

Previous studies have demonstrated that CAHS12 protein assembly is concentration-dependent, with higher concentrations facilitating the formation of extended networks and filamentous structures^65–67^. To investigate this phenomenon, we prepared two systems containing 10 and 20 CAHS12 protein models, respectively. The proteins were randomly positioned within cubic water boxes with a length of 40 nm. Each system was subjected to energy minimization, followed by 10 ps of NVT equilibration and 50 ns of NPT equilibration. Production simulations were then carried out for 1500 ns to allow for the emergence of protein aggregates, and the terminal 500 ns were subjected to analysis. To quantitatively assess protein assembly, we monitored the number of contacts formed between the N-terminal and/or C-terminal regions across the trajectories. A contact was defined as the COM distance between terminal regions falling below a 4 nm cutoff. Each data point represents an average contact number from the local 100 ns simulation window, and the uncertainty was computed using five-fold block averaging. The results indicate that higher protein concentration consistently yielded a greater number of terminal-region contacts, in agreement with experimental observations that elevated CAHS protein levels enhance the protective effects on GUV membranes.

#### Simulations of the CAHS12 Proteins and POPC Membrane

The CAHS12 protein is known to form extended protein networks that contribute to the structural stability of GUVs under dehydration^9,58^. It has also been speculated that CAHS12 interacts directly with lipid membranes, thereby enhancing its protective effects^68^. To explore these mechanisms, we performed large-scale coarse-grained simulations containing 40 CAHS12 proteins and a large patch of POPC membrane. Additionally, we included ΔC and ΔN, and ΔCΔN variants lacking, respectively, the C and N terminal domains to assess the respective contributions of the terminal regions to membrane association and network formation (**Supplementary Fig. 15, Supplementary Tables 1, 4, 5, 6**).

A small patch of POPC membrane patch consisting of 294 phospholipids was first constructed and solvated with 1 nm of water on each leaflet using packmol^69^, followed by a 4 ns equilibration in the NPT ensemble. Then, 25 such equilibrated membrane patches were tiled into a cuboidal simulation box to construct a larger membrane with a lateral dimension of 40 nm. The water layer thicknesses were then extended to 32 nm above and 5 nm below the membrane to accommodate protein insertion in the upper water bulk. This system was further equilibrated for 100 ns in the NPT ensemble. Subsequently, in 3 separate simulations, 40 wildtype CAHS12 proteins, ΔC, ΔN and ΔCΔN variants were randomly introduced into the bulk water above the membrane. The systems were then energy-minimized followed by a 50 ns of NPT equilibration. The production run was performed for 1.5 μs to allow the system to fully relax and undergo assembly. We acknowledge that simulation time is relatively short for these systems, but we observe the formation of stable aggregates that do not undergo substantial structural changes, at least over microsecond time scales, and anticipate the molecular behaviors observed in these simulations can be taken to be representative of the stable aggregates. Each system was prepared and simulated in three independent replicates, leading to a total of 12 simulations. Protein interaction networks were constructed using MDAnalysis^70,71^ and NetworkX^72^ using the final 500 ns of simulation trajectories. Each CAHS12 protein was represented as a node in the network, and edges were defined by inter-protein contacts, where a contact was considered present if any residue of one protein was within 0.4 nm of a residue from another protein. Edge weights were assigned as the inverse of the average COM distance between the two proteins. This network representation enabled characterization of the dynamic organization of protein assemblies, including connectivity, clustering behavior, and the contributions of individual proteins to the network topology. Key network metrics, including the number of clusters, average node degree (i.e., connectivity), and the size of the largest cluster, were extracted from three independent replicates for each system. Statistical significance between systems was performed using Welch’s t-test.

The contacts between CAHS12 proteins (and their terminal truncation variants) and the POPC membrane were assessed using the final 500 ns of the simulation trajectories. To define a proximal contact between a protein residue and the membrane, we first identified the local membrane region within a radius of 0.8 nm in the XY plane surrounding each protein residue. For each identified region, we evaluated the Z-coordinate of the residue relative to the surrounding lipid headgroups. A residue was classified as membrane-inserted if its Z-position fell within the vertical extent defined by the upper and lower boundaries of the lipid headgroup region. For each residue in the protein, the residue center of mass (COM) was projected onto the membrane plane, and all POPC NC3 headgroups within 8 Å of the COM position in the XY plane were selected. The Z-coordinates of these headgroups were then used to define a local membrane profile. The median of these Z-coordinates was used to separate the headgroups into upper and lower leaflet populations. The mean leaflet Z-positions were computed for both leaflets, and the membrane midplane was defined as the average of the upper and lower leaflet means. A residue was considered inserted if its COM Z-position lay between the midplane and the mean position of the nearest leaflet. This criterion ensured that only residues embedded within the membrane interface, rather than merely in proximity to it, were counted as forming meaningful membrane contacts. For the wild-type CAHS12 system, we further performed contact frequency analysis for each residue across distinct protein segments to identify key regions involved in protein-membrane association. Contact data were aggregated from three independent simulation replicates. The statistical comparisons between the wild-type and variant systems were conducted using Welch’s t-test to assess the significance of observed differences in contact patterns.

To mimic desiccation conditions, we performed water depletion simulations starting from one of the final structures from the previous simulations of full-length CAHS12 and its variants. The total number of water beads in the fully hydrated system was defined as 100%, and water content was reduced in 10% increments down to 10%, resulting in a series of nine dehydration states. For each dehydration level, the system was energy-minimized, followed by 20 ns of NVT equilibration and a 20 ns production run in the NPT ensemble. To evaluate how dehydration affects protein-membrane interactions, we quantified residue-level contacts between CAHS12 and the lipid bilayer across these dehydration windows. Specifically, we computed the number of protein residues within 0.5 nm of any lipid headgroup PO_4_ over the simulation trajectory. The results were averaged over time to obtain the mean number of contacting residues for each hydration level. This analysis revealed the progressive enrichment of protein-membrane contacts as hydration levels decreased, offering molecular insight into how CAHS12 enhances membrane stabilization under desiccating conditions.

#### Implementation Details

In all simulations, energy minimization was carried out using the steepest descent^73^ algorithm to eliminate forces larger than 1000 kJ/mol·nm^2^. For all simulations, the temperature was set to 300 K by velocity-rescale^74^ thermostat with a time constant of 2 ps. For NPT simulations, the pressure was maintained to be 1 bar by a c-rescale^75^ barostat with a time constant of 12 ps and a compressibility of 3.0×10^−4^ bar^-1^. The coupling scheme was isotropic for cubic boxes and semi-isotropic for simulations with POPC membranes^76^. The time step was set to 10 fs for equilibration and 20 fs for production, and Newton’s equations of motion were integrated by the leap-frog algorithm^77^. Lennard-Jones interactions were smoothly shifted to zero at a cutoff of 1.1 nm, and electrostatics were treated using the reaction field method with ε_rf_ = ∞ and ε_r_ = 15 as recommended for the non-polarizable Martini water model. All simulations were performed using the Gromacs 2022.4/2022.5 simulation suites^78–80^. The analyses of the simulation trajectories were performed using MDAnalysis^70,71^. ChimeraX^81^ was used to visualize the simulation snapshots. Plots were generated using matplotlib^82^ and seaborn^83^.

## Data and materials availability

Gromacs scripts for conducting coarse-grained molecular dynamics simulations of CAHS12 proteins and their interactions with a POPC lipid membrane, along with Jupyter notebooks for analyzing the simulation trajectories and generating plots, are available at https://github.com/Ferg-Lab/CAHS-Protein-Simulations.git. Experimental data are available upon request.

## Acknowledgements

We thank Takekazu Kunieda (University of Tokyo) for sharing the CAHS12 plasmid. We thank Alexandre Bisson (Brandeis University) and the Liu lab for helpful discussion.

## Author contributions

Conceptualization: Y.X., A.P.L. Methodology: Y.X., J.M., H.M., A.L.F., A.P.L. Investigation: Y.X., J.M., S.J.C., Y.J.L., A.P., A.J.H., D.Z., A.X. K.F.. Funding acquisition: A.L.F., A.P.L. Writing – original draft: Y.X., J.M., S.J.C., Y.J.L., A.P., A.L.F., A.P.L. Writing – review & editing: Y.X., J.M., S.J.C., H.M., Y.J.L., A.P., A.J.H., D.Z., A.X., K.F., S.A., A.L.F., A.P.L..

## Additional information

**Supplementary information** The online version contains supplementary material available at

**Correspondence and requests for materials** should be addressed to Andrew L. Ferguson or Allen P. Liu.

