## Supplementary Information for "Cytoplasmic Abundant Heat-Soluble Proteins from Tardigrades Protect Synthetic Cells Under Stress"


**The PDF file includes:**

Supplementary Method
Supplementary Figures 1-28
Supplementary Tables 1-7
References

### 1. Supplementary Method

#### Stability, thermal tolerance, and osmotic stress assays

To assess long-term stability under ambient and desiccated conditions, giant unilamellar vesicles (GUVs) encapsulating CAHS12-GFP and Cy5 dye (300 mM sucrose interior, 300 mM glucose exterior) were prepared by phase transfer and stored in the dark at room temperature (22–25 °C) for up to 74 days. At designated time points (0.5, 5, 14, 30, and 74 days), vesicles were imaged using confocal fluorescence microscopy. Vesicle morphology and diameter were quantified using FIJI (ImageJ, NIH) based on Cy5 and GFP signal. Vesicles were considered intact if they retained spherical shape and luminal fluorescence. Protein-free GUVs and those encapsulating CAHS12 ΔN served as negative controls.

To examine the effects of dehydration duration, GUVs were deposited in 96-well plates and subjected to vacuum desiccation at room temperature ($10^{-2}$ mbar) for varying durations (0.5–14 days). After rehydration with ultrapure water, vesicle integrity was assessed by confocal imaging. Circularity and size distributions were extracted using custom macros in ImageJ. Vesicle populations were defined as structurally preserved if >85% of vesicles retained a circularity >0.75.

To assess thermal stability, GUV samples were prepared as described above and transferred into 0.2 mL low-retention PCR tubes (Axygen). The tubes were sealed to minimize water exchange with the environment and placed in a thermal cycler (Bio-Rad T100) programmed to maintain constant temperatures of 25°C, 37°C, 45°C, 60°C, or 80°C for the indicated durations (typically 2 hours to 5 days). After incubation, samples were allowed to cool to room temperature. GUV structural integrity was subsequently assessed by fluorescence microscopy to evaluate vesicle morphology, size distribution, and leakage. Vesicle survival was defined as the number of vesicles in 3 random fields of view relative to day 0.

Osmotic stress response was tested by mixing equal volumes (5 µL each) of GUV suspension and hyperosmotic glucose solution (1200 mOsm/kg, prepared in ultrapure water) to generate an external osmotic pressure 2.5× that of the GUV lumen (300 mOsm/kg). Vesicles were imaged at 0, 5, and 40 minutes post-mixing using fluorescence microscopy. Structural integrity was assessed by quantifying vesicle diameter and circularity. For all experiments, at least 100 vesicles per condition were analyzed. Results represent means ± s.d. from three independent replicates.

#### Fluorescence microscopy and image analysis

Fluorescence images were acquired using an Olympus IX-81 inverted microscope equipped with a spinning disk confocal system (Yokogawa CSU-X1) and a solid-state laser module (Solamere Technology, controlled via National Instruments DAQmx). The system was configured with an iXON3 electron-multiplying CCD camera (Andor Technology) and a 40×/1.4 NA Plan-Apochromat oil immersion objective (Olympus) to capture high-resolution images of GUVs under hydrated and dehydrated conditions. A 60x/1.42 Plan-Apo N oil immersion objective (Olympus) lens was used to image SpyTag GUVs and SpyCatcher cells. Image acquisition parameters were controlled using MetaMorph software (Molecular Devices).

Fluorescence signals from CAHS12-GFP, rhodamine-PE (Rhod-PE), mCherry, and Cy5 were sequentially captured using 488 nm, 561 nm, 561 nm, and 647 nm laser excitation, respectively. Emission was collected using bandpass filters optimized for each fluorophore. Typical exposure times were 200 ms for 488 nm excitation and 400 ms for both 561 nm and 647 nm excitation channels. Z-stack imaging was performed with step sizes of 0.2 µm where needed to confirm uniform encapsulation, internal distribution of protein, and vesicle morphology.

Acquired images were processed using Fiji (ImageJ, NIH) for quantitative analysis of vesicle diameter[1], circularity, and fluorescence intensity. Vesicle boundaries were identified using automated thresholding followed by particle analysis to extract geometric parameters, with manual curation to exclude aggregates or artifacts. Only vesicles with diameters ⩾ 2 µm and in focus (determined by maximal fluorescence cross-section) were included in downstream analyses. Circularity was computed using the standard Fiji metric $4\pi(area/perimeter^2)$ to assess vesicle integrity, with values closer to 1 indicating higher sphericity.

For quantitative fluorescence analysis, regions of interest (ROIs) were drawn around individual vesicles to extract mean fluorescence intensity within the vesicle lumen and at the membrane, where applicable. Background fluorescence was subtracted using mean pixel values from nearby vesicle-free regions. Statistical data were compiled from at least three independent experiments, each with n > 100 vesicles per condition, to ensure reproducibility across preparations and conditions.

**Micropipette aspiration.**

Micropipettes were pulled from standard borosilicate glass capillaries (World Precision Instruments) using a P-87 pipette puller (Sutter Instruments) and fire-polished with a microforge (MF-83, Narishige) to yield an inner diameter of 8–10 μm. Micropipettes were mounted in a holder and connected via a continuous fluidic line to a filling syringe and a pressure reservoir through a three-way valve. The reservoir was pneumatically coupled to a high-speed pressure clamp (HSPC, ALA Scientific Instruments), allowing precise control of aspiration pressure.

The filling syringe contained a glucose solution osmotically matched to the external solution of the GUVs and was used to fill both the micropipette and the reservoir. All air bubbles were carefully eliminated from the system, as their presence compromises pressure stability during aspiration measurements. Using a micromanipulator, the micropipette was positioned in close proximity to a target GUV. Once the GUV was aligned, controlled negative pressure was applied via the pressure transducer to induce membrane aspiration into the micropipette. The applied suction pressure $\Delta P$ generates a uniform membrane tension $\tau$ in the vesicle membrane. The membrane tension is determined by the geometry of the system and depends on the inner radius of the micropipette ($R_p$) and the instantaneous radius of the vesicle exterior to the micropipette ($R_{GUV}$), according to:

$$\tau = \frac{(R_p \Delta P)}{\left(2\left(1 - \frac{R_p}{R_{GUV}}\right)\right)}$$

The apparent membrane area strain (*ΔA/A*) was calculated from the aspirated tongue length L, the pipette radius ($R_p$), and the vesicle geometry. The change in membrane area was normalized by the initial vesicle surface area, defined by the unstressed vesicle radius ($R_{0,GUV}$), yielding:

$$\frac{\Delta A}{A} = \frac{\left(2\pi\, R_p \Delta L\, \left(1 - \frac{R_p}{R_{GUV}}\right)\right)}{\left(4\pi\, R_{0,GUV}^2\right)}$$

To determine the membrane area stretch modulus ($K_a$), the membrane tension $\tau$ was plotted as a function of the direct area expansion (*ΔA/A*). The resulting linear relationship was fit by least-squares regression, and the slope of this plot corresponds to the area stretch modulus $K_a$[2,3]

**Statistical analysis**

Unless otherwise noted, data are presented as mean ± standard deviation (s.d.). Statistical significance between groups was determined using Welch's two-tailed *t*-test to account for unequal variances. For vesicle-based assays—including measurements of circularity, survival rate, and interfacial tension (IFT) were analyzed per condition across at least three independent biological replicates. Sample sizes and replicate numbers are specified in figure legends. All image-based quantifications were performed using ImageJ (Fiji).

### 2. Supplementary Figures

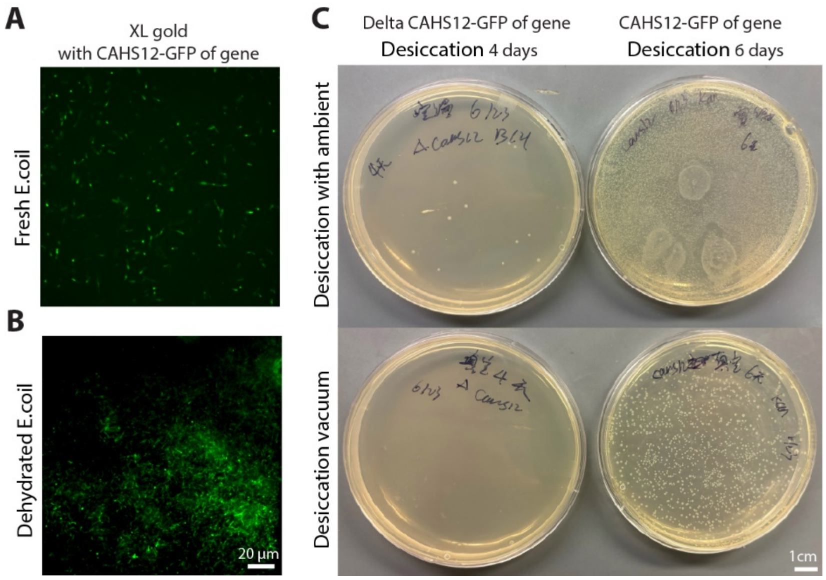


**Supplementary Fig. 1. CAHS12-GFP enhances *E. coli* survival after desiccation under ambient and vacuum desiccation conditions.** (**A**) Fluorescence microscopy image of freshly prepared XL-10 gold *E. coli* cells expressing CAHS12-GFP. (**B**) Fluorescence image of *E. coli* expressing CAHS12-GFP after dehydration for 6 days. Scale bar, 20 µm. (**C**) Colony formation assay showing viability of *E. coli* expressing CAHS12-GFP after desiccation under ambient conditions (top) and vacuum (bottom). Plates on the left show wildtype *E. coli* after 4 days of desiccation, with minimal colony formation. Plates on the right show cells expressing CAHS12-GFP after 6 days of desiccation before plating, exhibiting robust colony recovery under both ambient and vacuum conditions. This result is consistent with prior studies reporting increased bacterial desiccation tolerance upon heterologous expression of CAHS-family proteins[4–6]. Scale bar, 1 cm.

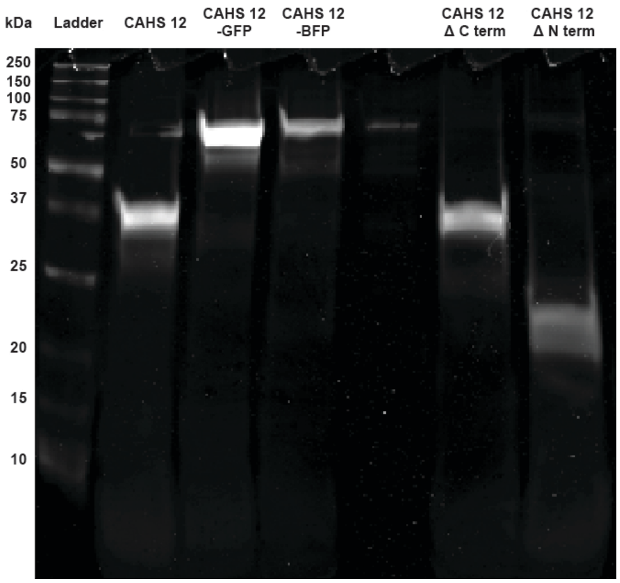


**Supplementary Fig. 2. SDS–PAGE analysis of purified CAHS12 and truncation variants.** Coomassie-stained SDS–polyacrylamide gel showing full-length CAHS12 and domain-deletion variants after Ni-NTA affinity purification. Lanes: molecular weight marker (kDa); 1, CAHS12; 2, CAHS12-GFP; 3, CAHS12-BFP; 4, CAHS12 ΔC; 5, CAHS12 ΔN. All proteins were expressed in *E. coli* BL21(DE3) and purified via $His_6$-tag affinity chromatography followed by buffer exchange into Tris-based storage buffer (pH 8.0). Predicted molecular weights: CAHS12, ~32 kDa; CAHS12-GFP, ~60 kDa; CAHS12-BFP, ~60 kDa; ΔC, ~29 kDa; and ΔN, ~16 kDa.

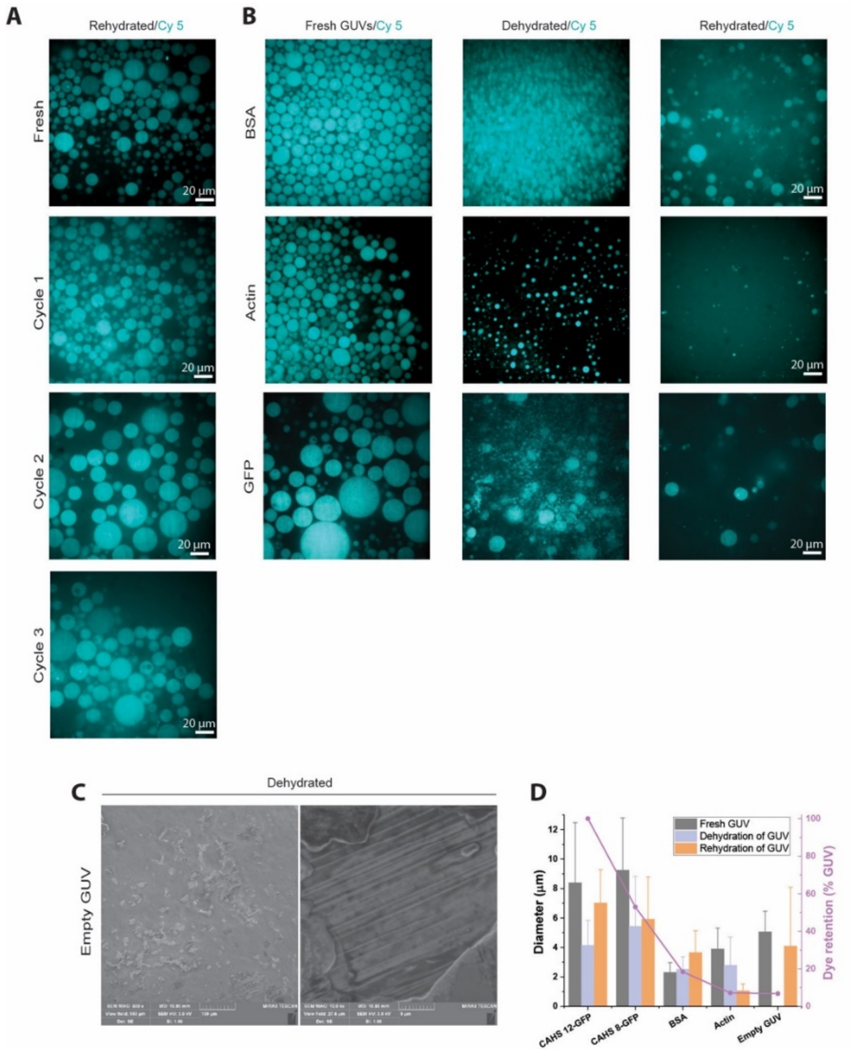


**Supplementary Fig. 3. GUVs lacking CAHS12 fail to retain structural integrity after dehydration–rehydration.** (**A**) Representative fluorescence microscopy images of GUVs following three consecutive dehydration–rehydration cycles. After each dehydration step (vacuum desiccation at ~31 mbar for ⩾12 h), samples were rehydrated by adding ultrapure water and incubated at room temperature for 10 min to allow complete rehydration before imaging or the initiation of the subsequent cycle. Vesicle morphology and integrity were assessed after each rehydration. Some vesicles appeared larger after the second dehydration-rehydration cycle, for reasons we do not fully understand. (**B**) Representative fluorescence microscopy images of GUVs encapsulating 280 μM bovine serum albumin (BSA), 10 μM actin, or 280 μM GFP under fresh, dehydrated, and rehydrated conditions. Cy5 dye was co-encapsulated as a membrane integrity reporter; dye retention indicates vesicle stability, while loss of fluorescence reflects membrane rupture or leakage. (**C**) Scanning electron microscopy (SEM) images of empty GUVs after vacuum dehydration, showing morphological collapse and wrinkling. left: low magnification overview; right: high magnification of individual vesicles. Scale bars, 100 μm (top) and 5 μm (bottom). (**D**) Quantitative comparison of GUV size (mean diameter) and dye retention fraction after dehydration–rehydration for vesicles containing different protein cargos. Only CAHS12-stabilized GUVs (n=198-220) preserved size and structural integrity, whereas BSA- (n= 52-283), actin-loaded vesicles (n= 17-237), and empty GUVs (n=15-221) exhibited significant shrinkage and failure to rehydrate. Data represent mean ± s.d.; survival is defined as retention of the number of GUVs.

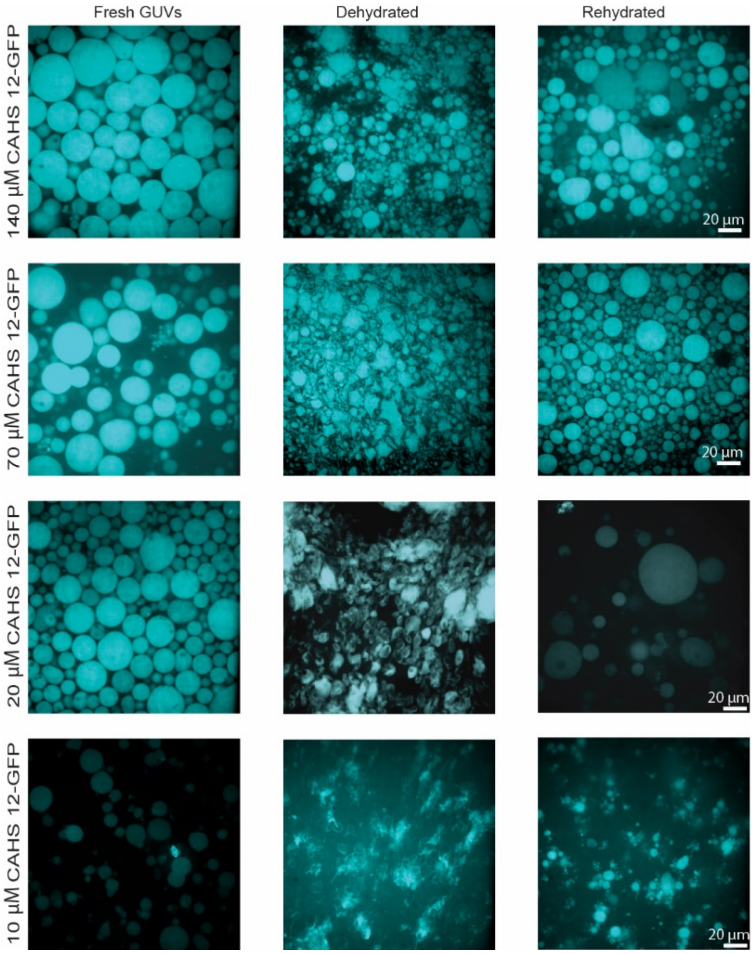


**Supplementary Fig. 4. Dose-dependent protection of synthetic cells by CAHS12-GFP during dehydration–rehydration.** Fluorescence microscopy images of GUVs encapsulating varying concentrations of CAHS12-GFP (10, 20, 70, and 140 µM) under fresh, dehydrated, and rehydrated conditions. Cy5 dye was co-encapsulated as a fluorescent reporter of membrane integrity; retention of Cy5 fluorescence indicates preservation of vesicle structure, whereas dye loss signifies membrane rupture or leakage. Increasing concentrations of CAHS12-GFP correlated with improved structural preservation and dye retention following dehydration. Scale bars, 20 µm. Representative images shown from ≥3 independent experiments.

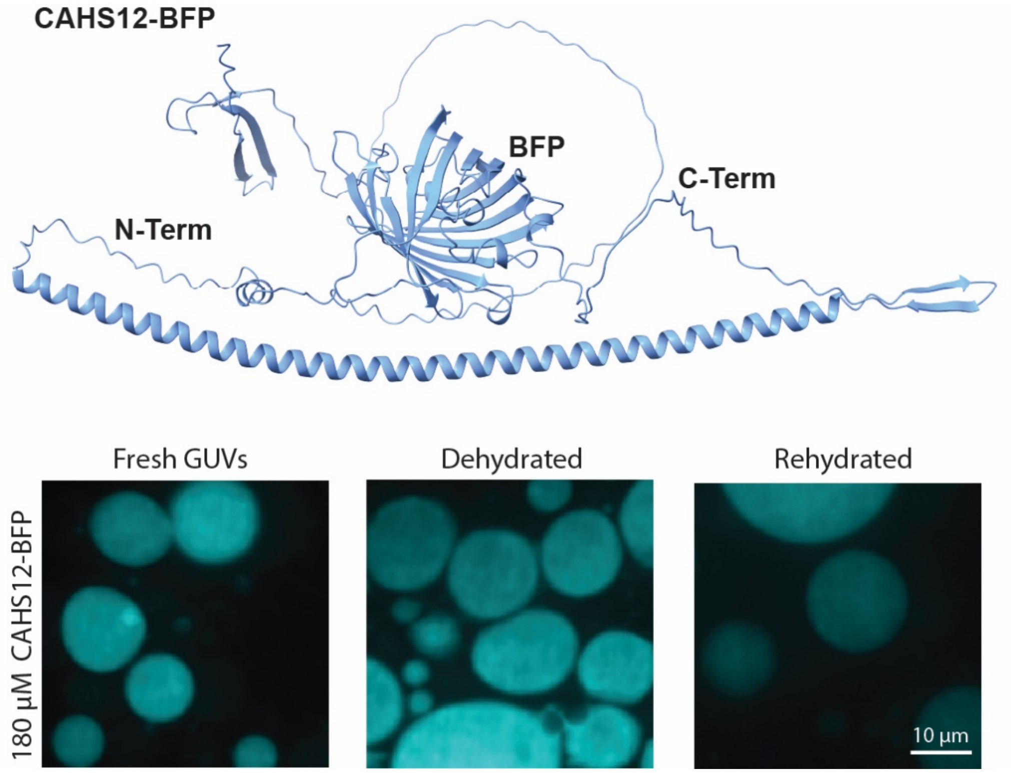


**Supplementary Fig. 5. Structural preservation of synthetic cells by CAHS12-BFP during dehydration–rehydration.** (Top) Proposed model of CAHS12-BFP, secondary structures of CAHS12-BFP were predicted using AlphaFold 3, and molecular rendering of CAHS12 protein was constructed using ChimeraX[7]. (Bottom) Fluorescence microscopy images of GUVs encapsulating 180 µM CAHS12-BFP fusion protein under fresh, dehydrated, and rehydrated conditions. Cy5 dye was co-encapsulated as a membrane integrity reporter to assess vesicle stability; retention of Cy5 fluorescence indicates preserved membrane structure, while fluorescence loss reflects leakage or rupture. CAHS12-BFP localized within vesicles and conferred robust protection against dehydration-induced damage. Scale bars, 20 µm. Images representative of ≥3 independent experiments.

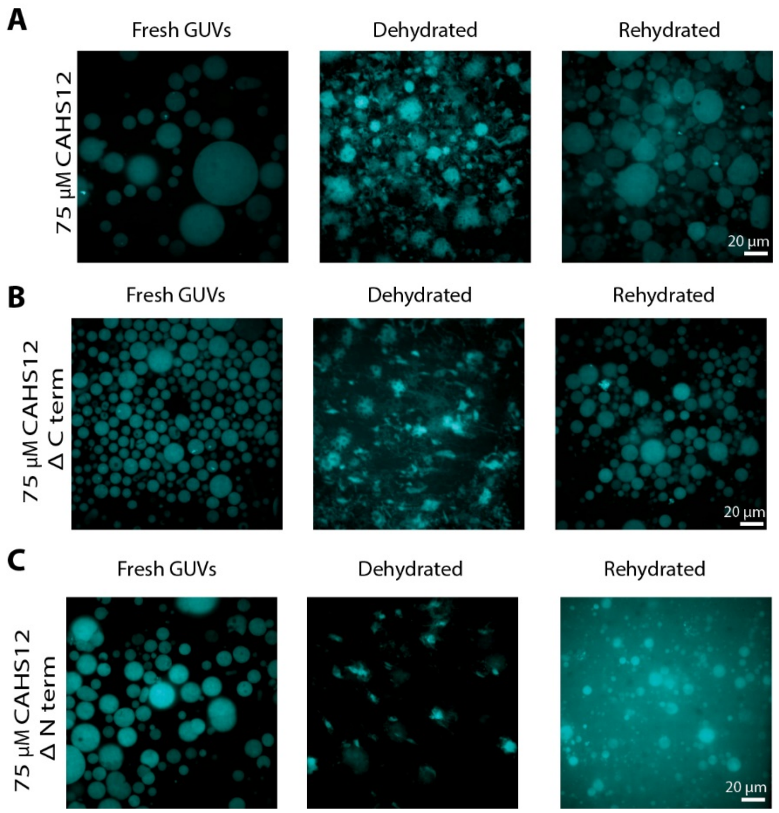


**Supplementary Fig. 6. Functional contributions of CAHS12 termini to desiccation protection in GUVs.** Fluorescence microscopy images of GUVs encapsulating 75 µM full-length CAHS12 (**A**), 75 µM C-terminal truncation (**B**), or 75 µM N-terminal truncation (**C**) under fresh, dehydrated, and rehydrated conditions. Cy5 dye was co-encapsulated as a membrane integrity reporter. Full-length CAHS12 and ΔC maintained vesicle integrity following dehydration and rehydration, while ΔN failed to prevent membrane collapse or dye leakage. Representative images from ≥3 independent experiments.

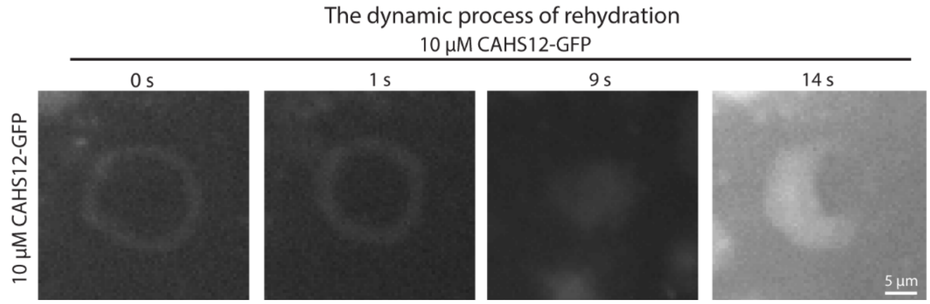

**Supplementary Fig. 7.** The dynamic process of dehydration and rehydration of GUVs containing 10 μM of CAHS12-GFP. Scale bar, 5 μm.

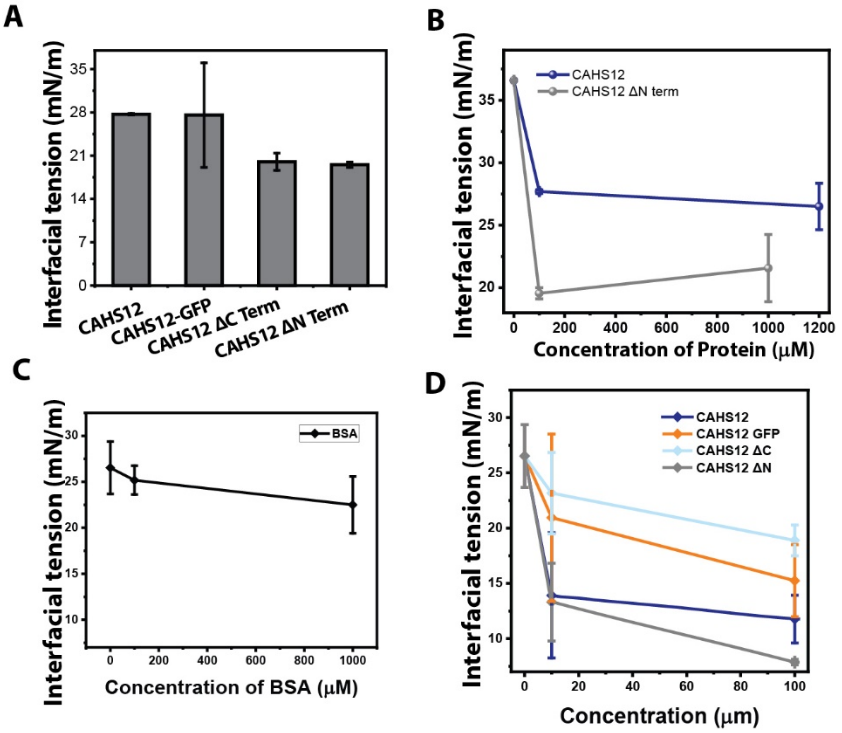


**Supplementary Fig. 8. CAHS12 modulates interfacial tension in a construct- and concentration-dependent manner.** (**A**) Quantification of interfacial tension (IFT) of aqueous droplets containing 0.1 mM full-length CAHS12, CAHS12-GFP, CAHS12 ΔC, or CAHS12 ΔN in mineral oil lacking POPC, measured using a hanging drop tensiometer. (**B**) Dose-dependent IFT profiles for CAHS12 and its truncation mutants (ΔN) across a concentration range of 0–1 mM. Data represent mean ± s.d. from three independent measurements per condition. (**C**) IFT as a function of concentration for BSA. (**D**) IFT as a function of concentration for CAHS12, CAHS12 GFP, CAHS12 ΔC, and CAHS12 ΔN.

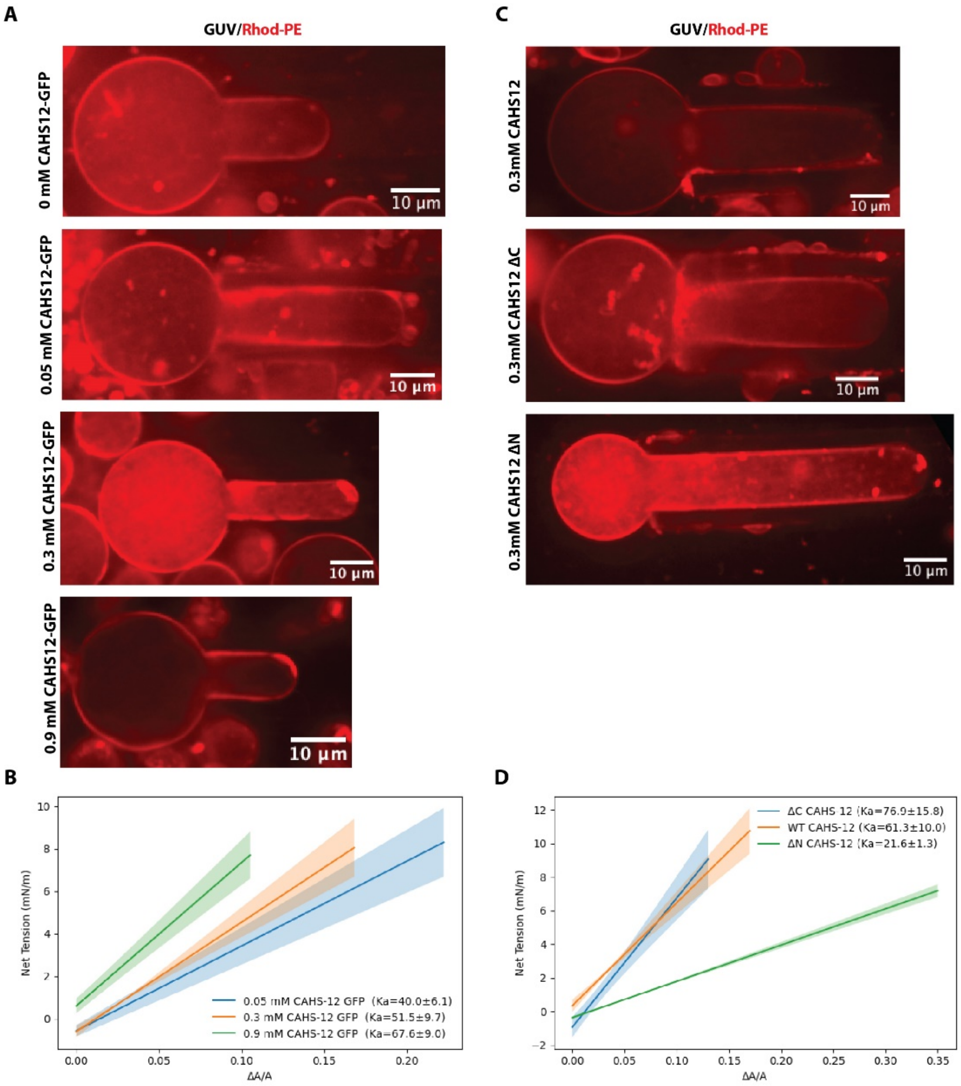


**Supplementary Fig. 9. Micropipette aspiration reveals construct- and concentration-dependent mechanical stabilization of GUVs by CAHS12.** Representative fluorescence microscopy images of Rhod-PE–labeled GUVs (red) subjected to micropipette aspiration at an applied aspiration pressure of 2660 Pa. (**A**) GUVs encapsulating increasing concentrations of CAHS12–GFP (0, 0.05, 0.3, and 0.9 mM) show a concentration-dependent increase in resistance to deformation under applied membrane tension. Empty GUVs exhibit relatively short aspiration lengths, likely due to higher initial membrane tension limiting available excess area. At low CAHS12 concentration (0.05 mM), partial membrane adsorption transiently increases effective deformability, whereas at higher concentrations (≥0.3 mM), interfacial protein networks mechanically stabilize the membrane, reducing aspiration length. **(B)** Concentration-dependent modulation of membrane mechanics by CAHS-12–GFP. Linear fits to net membrane tension versus areal strain are shown for GUVs containing increasing concentrations of CAHS-12–GFP (0.05, 0.3, and 0.9 mM). Solid lines represent the mean linear fit across three independent replicates for each concentration, with shaded regions indicating plus or minus one standard deviation across replicate fits. The apparent area expansion modulus ($K_a$) increases with CAHS-12–GFP concentration, from 40.0 ± 6.1 mN/m at 0.05 mM to 51.5 ± 9.7 mN/m at 0.3 mM and 67.6 ± 9.0 mN/m at 0.9 mM, indicating enhanced resistance to aspiration-induced deformation. Linear fits for individual replicates yielded R-squared values ranging from 0.84 to 0.98 (0.05 mM), 0.91

to 0.97 (0.3 mM), and 0.94 to 0.99 (0.9 mM). N = 3 independent experiments. (**C**) GUVs encapsulating full-length CAHS12, CAHS12 ΔC, or CAHS12 ΔN at the same protein concentration. GUVs containing full-length CAHS12 or CAHS12 ΔC largely retain a spherical morphology during aspiration, whereas GUVs containing CAHS12 ΔN exhibit pronounced elongation, indicating reduced mechanical stability. **(D)** Quantification of $K_a$ from linear fits to net membrane tension versus areal strain obtained by micropipette aspiration of GUVs containing CAHS-12 variants. Solid lines represent the mean linear fit across three independent replicates for each condition, with shaded regions indicating plus or minus one standard deviation across replicate fits. GUVs containing the C-terminal deletion mutant (ΔC CAHS-12) exhibit the highest apparent Ka (76.9 ± 15.8 mN/m), followed by wild-type CAHS-12 (61.3 ± 10.0 mN/m), whereas the N-terminal deletion mutant (ΔN CAHS-12) shows a substantially reduced modulus (21.6 ± 1.3 mN/m). Linear fits for individual replicates showed strong agreement, with R-squared values ranging from 0.85 to 0.97 (ΔC), 0.96 to 0.99 (WT), and 0.82 to 0.96 (ΔN).

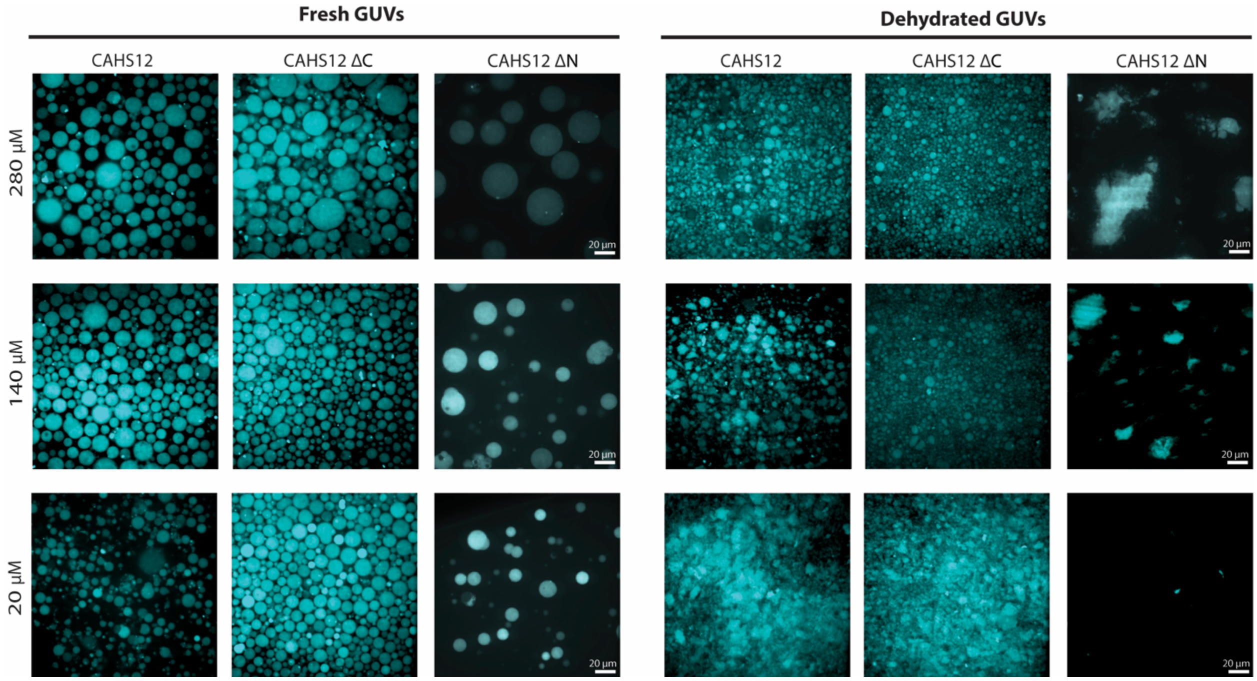


**Supplementary Fig. 10. Concentration-dependent protection of GUVs by CAHS12 and its truncation variants.** Fluorescence microscopy images of GUVs encapsulating full-length CAHS12, CAHS12 ΔC, or CAHS12 ΔN at increasing concentrations (20, 140, and 280 μM) under fresh and dehydrated conditions. Vesicle morphology and dye retention were used to assess membrane integrity. Full-length CAHS12 and CAHS12 ΔC exhibited concentration-dependent protection against desiccation-induced collapse. While CAHS12 ΔN showed minimal protective capacity at all tested concentrations. Scale bars, 20 μm. Images representative of ≥3 independent experiments.

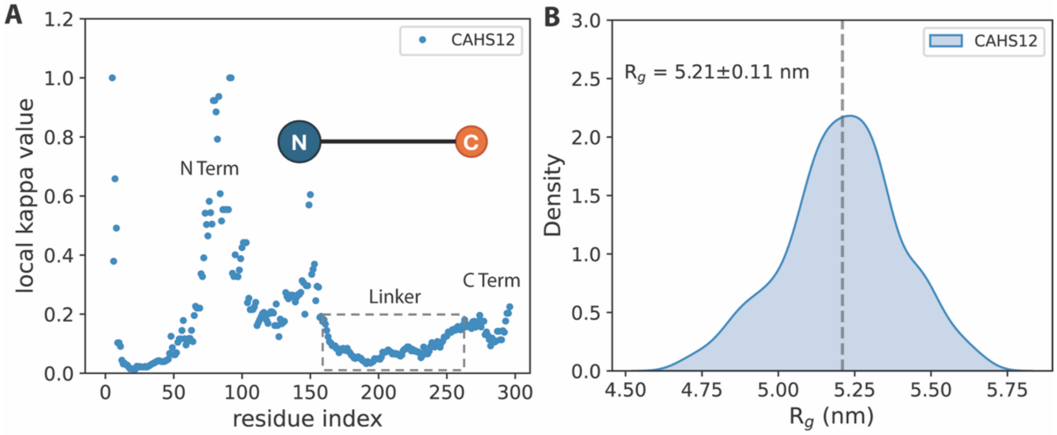


**Supplementary Fig. 11**. **Structural characterization of a single simulated CAHS12 protein.** (**A**) Local κ values along the CAHS12 sequence – a measure of charge patterning within a protein sequence, with high values indicative of disordered conformations – were calculated using a sliding window of 50 residues. The inset illustrates a cartoon of the CAHS12 protein, highlighting its dumbbell-like architecture comprising a rigid linker region that exists as an extended α-helix flanked by N and C termini that exist as random coils. (**B**) Kernel density estimation of the radius of gyration (Rg) distribution for a single CAHS12 protein obtained from 200 ns simulation. The dashed line indicates the average Rg values for a disordered coil-helix-coil topology.

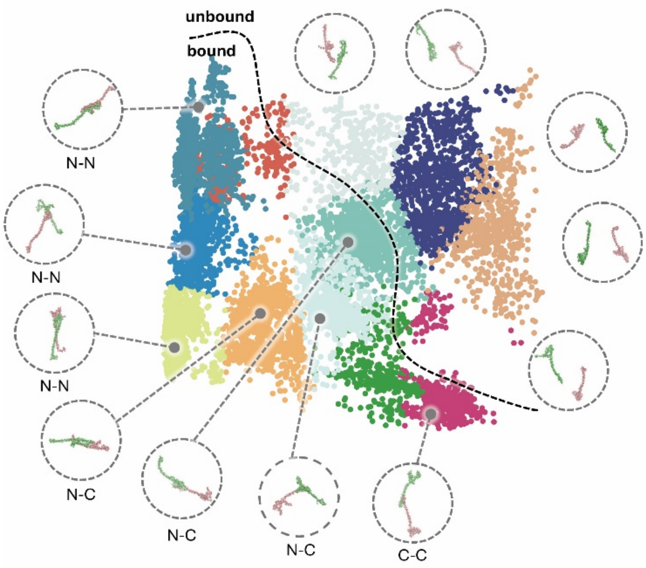


**Supplementary Fig. 12. Principal components analysis (PCA) of CAHS12 dimer conformations from unbiased simulations**. Each point represents a simulation frame projected onto the first two PCs derived from inter-protein features. Colors correspond to distinct binding modes identified via k-means clustering. The dashed line denotes the boundary between bound and unbound dimer states. Protein structures corresponding to representative binding modes were rendered using Visual Molecular Dynamics (VMD)[8].

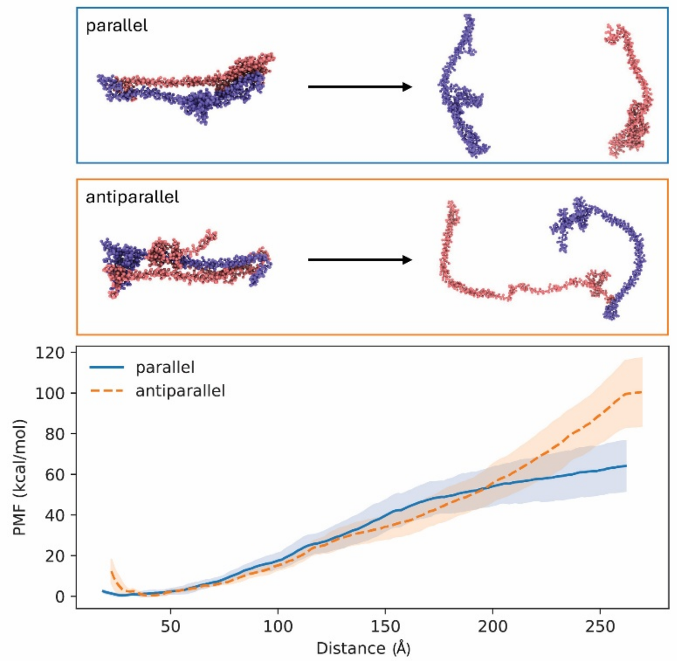


**Supplementary Fig. 13. Free energy calculations of dissociation of CAHS dimers from parallel and antiparallel conformations.** The upper panel shows representative initial bound structures and the corresponding final dissociated states. The lower panel reports the free energy profiles along the dissociation coordinate. Fifty window simulations with an interval of 0.5 nm were extracted and simulated using GROMACS and PLUMED[9]. The free energy profiles were computed using WHAM from the final 20 ns simulations[10]. The shaded areas present the uncertainty of the free energy profiles obtained from four-fold block analysis.

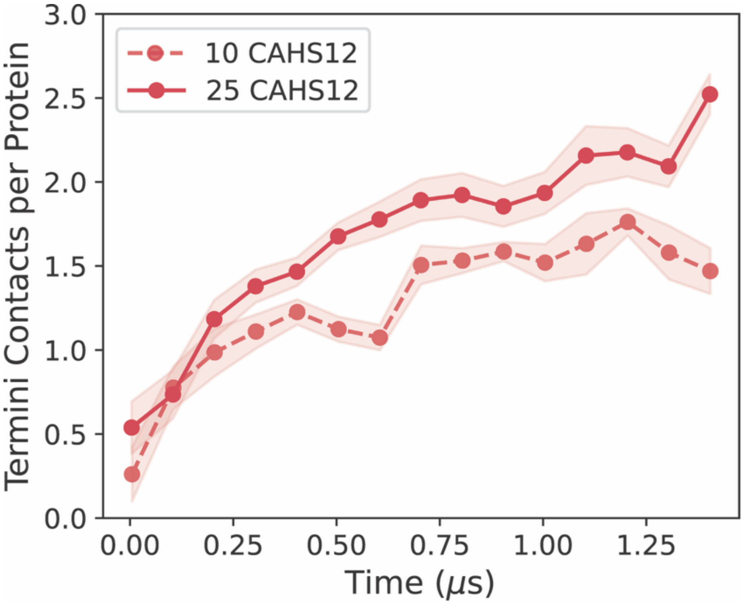


**Supplementary Fig. 14. Terminal-region contact formation of CAHS12 proteins at different concentrations.** The number of termini-termini contacts per protein is plotted over time for systems containing either 10 or 25 CAHS12 proteins. Contacts are defined based on center-of-mass distances between N- and/or C-terminal regions below a 4 nm cutoff. Shaded areas represent uncertainty obtained by five-fold block averaging in a 100 ns window.

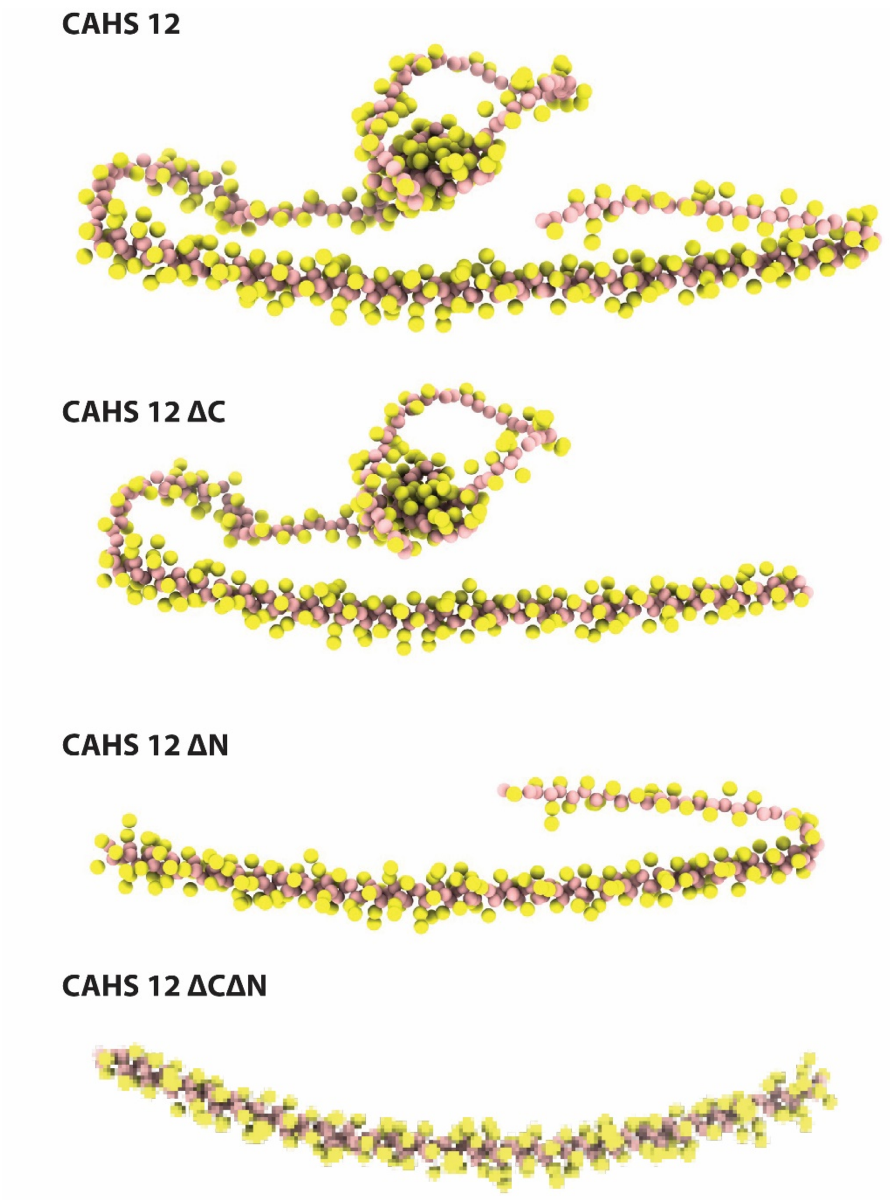


**Supplementary Fig. 15. Coarse-grained models of the CAHS12 protein and its terminal truncation variants (CAHS12 ΔC, CAHS12 ΔN and CAHS12 ΔCΔN).** The pink beads represent protein backbone while yellow beads represent side chains of residues. The models illustrate the overall architecture of the wild-type and truncated proteins used in simulations. Secondary structures were predicted using AlphaFold 3 and assigned during coarse graining using define secondary structure of proteins (DSSP) algorithm. The structures were visualized by ChimeraX[7].

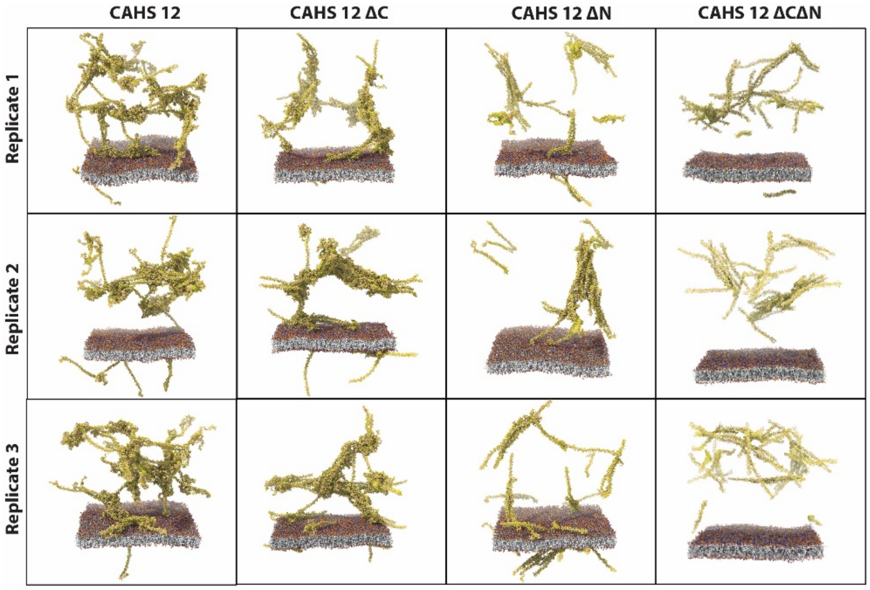


**Supplementary Fig. 16. Simulations of CAHS12 proteins and POPC membranes.** Final snapshots from each simulation were visualized using ChimeraX[7]. Three independent replicates were performed for each system, all of which yielded consistent phenomenological observations regarding protein assembly and membrane association.

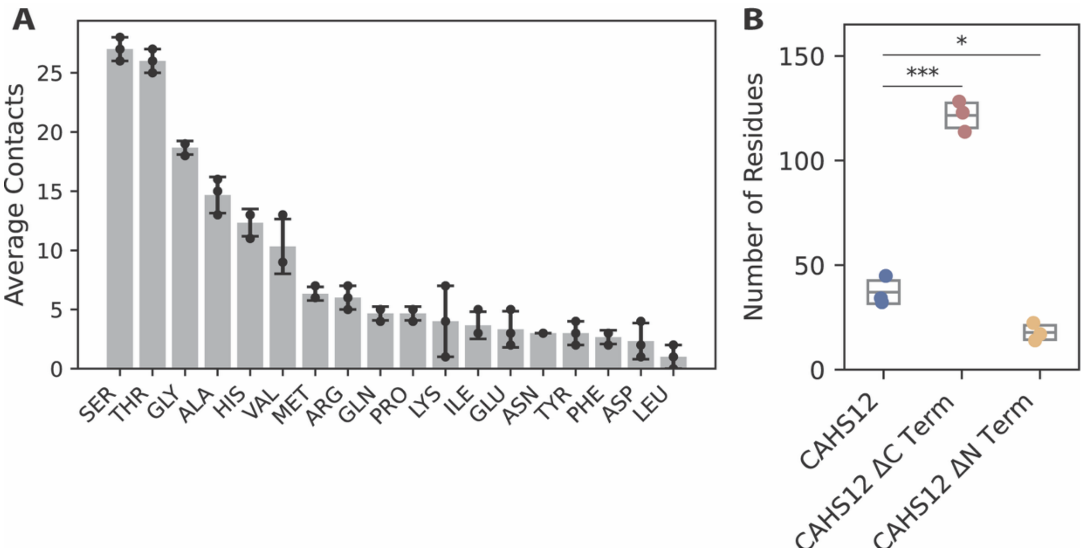


**Supplementary Fig. 17. Analysis of CAHS12-membrane contacts and the role of terminal regions in membrane association.** (**A**) Average number of membrane contacts per residue type observed in wild-type CAHS12 over the final 500 ns of simulation. Error bars represent standard deviations across three independent replicates. Polar and small residues such as Ser, Thr, and Gly exhibit the highest contact frequencies, suggesting a preferential interface with the lipid surface. (**B**) Number of residues in contact with the POPC membrane for CAHS12 and its ΔC and ΔN variants. Box plots show distributions across three replicates. Statistical significance was assessed using Welch's t-test, indicating a prominent role of the terminal regions, especially the N-terminus, in mediating membrane association. *, $p < 0.05$; **, $p < 0.01$; ***, $p < 0.001$.

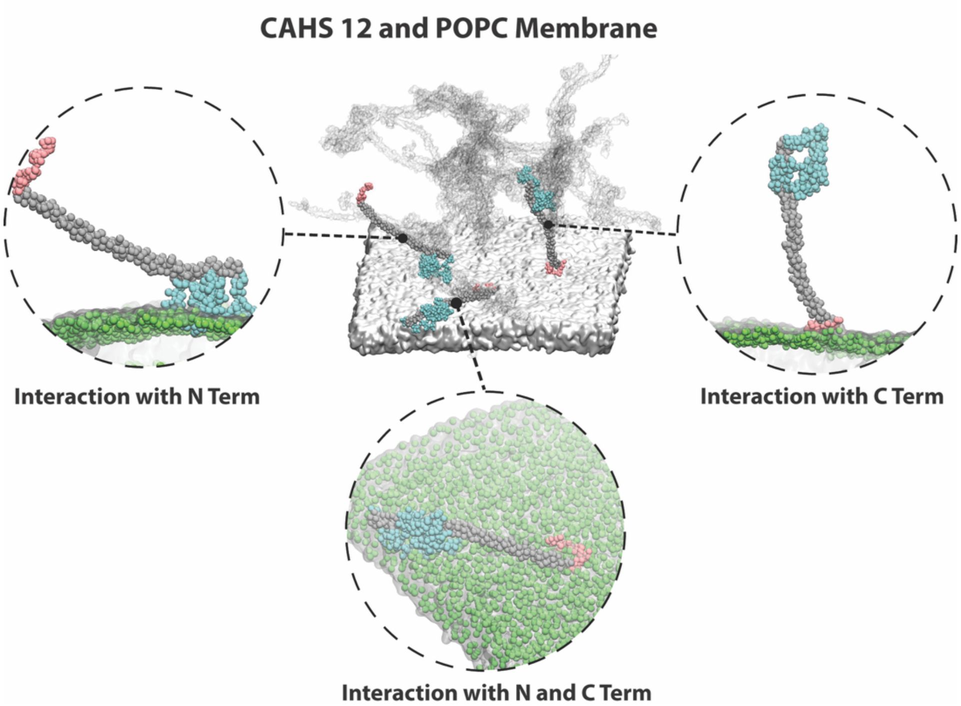


**Supplementary Fig. 18. Distinct modes of CAHS12 protein interaction with the POPC membrane.** Representative snapshots from coarse-grained simulations illustrating three major modes of membrane interaction by CAHS12 proteins. The central panel shows an overview of the protein-membrane system, highlighting several proteins engaging with the membrane via different domains. Insets depict zoomed-in views of proteins interacting through the N-terminal region (left), C-terminal region (right), and both N- and C-terminal regions (bottom). Linker regions are shown in gray, N-terminal residues in red, C-terminal residues in blue, and the POPC membrane in green (headgroups) and light gray (tails).

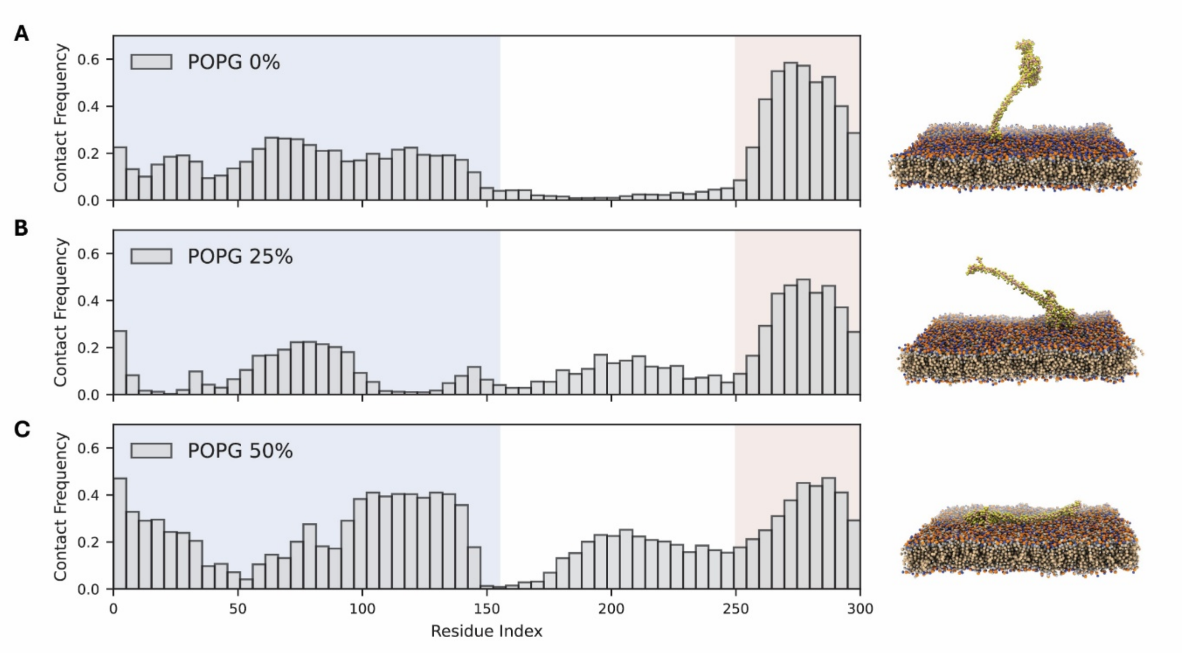


**Supplementary Fig. 19**. Membrane composition modulates CAHS12-membrane interactions. The residue level contact frequency of CAHS12 with POPC bilayers containing 0% (**A**), 25% (**B**), and 50% (**C**) POPG, along with their representative snapshots of the corresponding simulations. An increasing percentage in the POPG composition corresponds to increasing membrane surface charge and an observed increase in interactions between the linker region and the membrane. A contact is defined when the minimum distance between any residue atom and a lipid phosphate bead is less than 1 nm, and the contact frequency is reported as the fraction of simulation time each residue spends in contact with the membrane. Data were obtained from three independent simulations totaling 5 μs for each membrane composition and concatenated for analysis. To reduce local fluctuations and highlight regional trends along the sequence, contact frequencies are averaged over sliding windows of five residues, with each bar representing the mean value within a window. The simulation snapshots were rendered using ChimeraX[7].

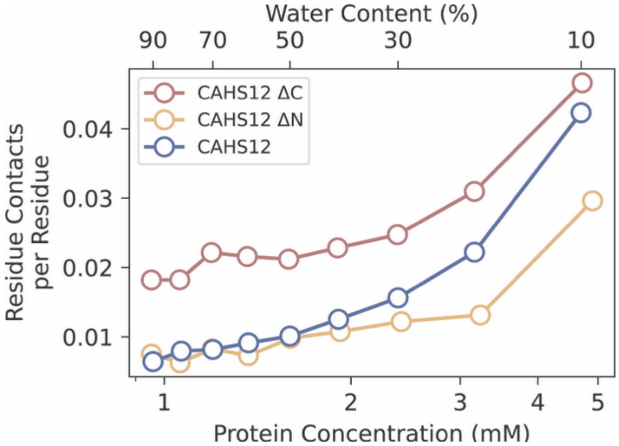


**Supplementary Fig. 20. Membrane contacts of CAHS12 and its variants with the POPC membrane under simulated dehydration.** Residue contacts with the POPC membrane, normalized per residue, are plotted as a function of protein concentration and water content for CAHS12, ΔC, and ΔN variants.

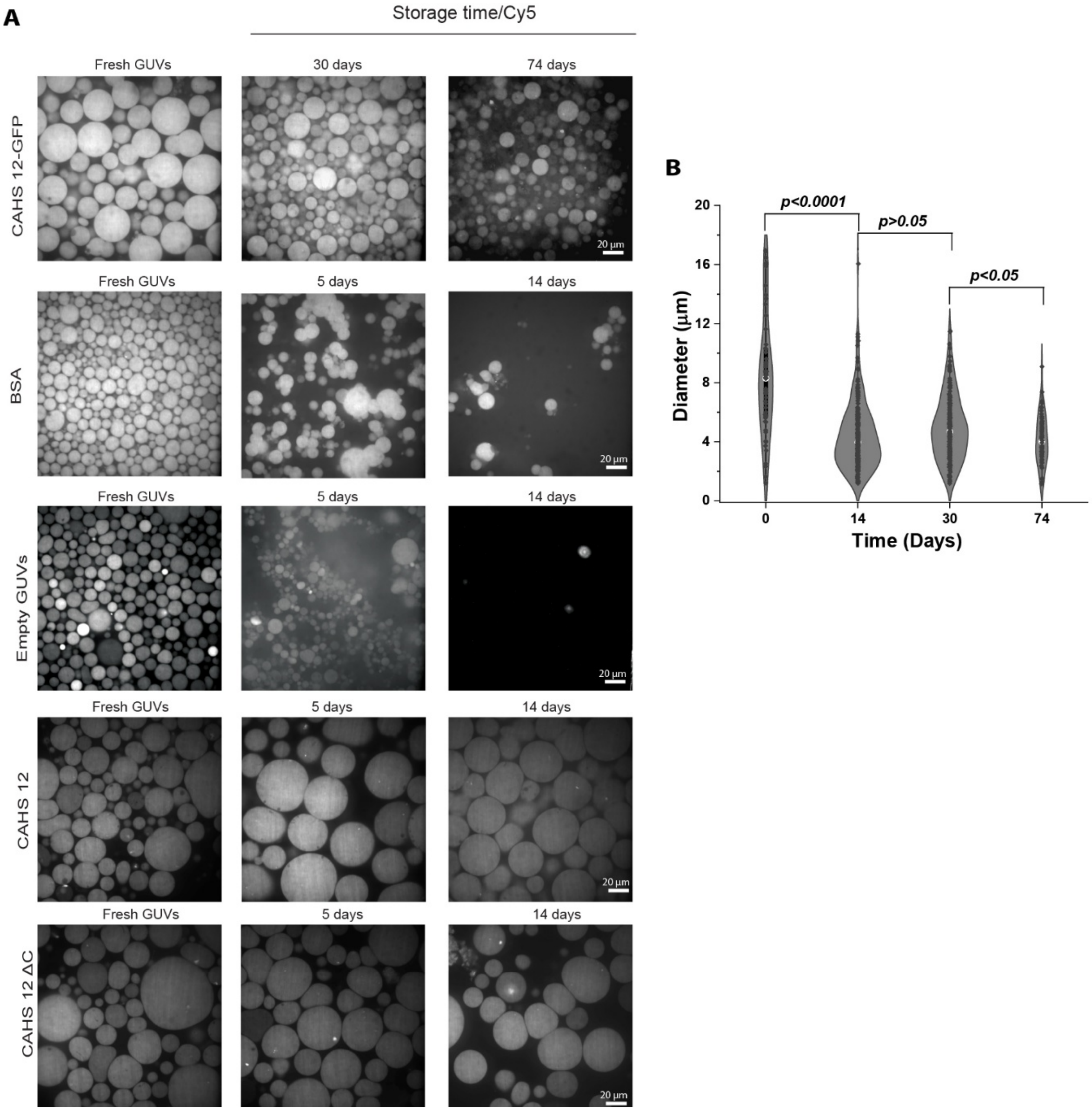


**Supplementary Fig. 21. Long-term ambient storage stability of empty GUVs and GUVs encapsulating CAHS12-GFP, BSA, CAHS12, or CAHS12 ΔC.** (**A**) Confocal fluorescence microscopy images of GUVs co-encapsulating Cy5 dye and either 280 μM CAHS12-GFP, BSA CAHS12, or CAHS12 ΔC, imaged after storage under ambient conditions. (**B**) Quantification of the diameter of CAHS12-containing GUVs following incubation for different numbers of days.

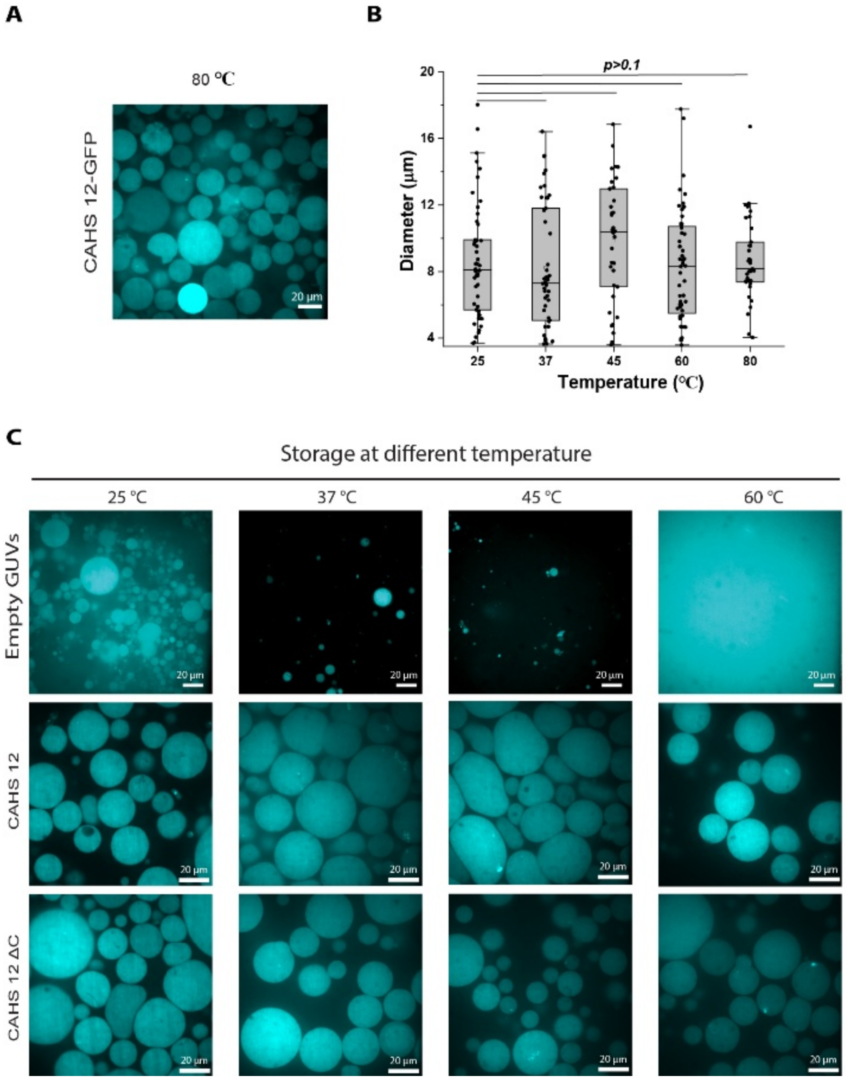


**Supplementary Fig. 22. Thermal stability and morphological response of GUVs under heat stress.** (**A**) Representative confocal fluorescence micrographs of GUVs containing CAHS12-GFP after incubation at 80 °C for 2 hours. (**B**) Quantification of GUV diameter following incubation at different temperatures (25 °C for 5 days, 37 °C for 2 days, 45 °C for 2 days, 60 °C for 2 hours, and 80 °C for 2 hours). Data represent mean ± s.d. of >100 vesicles per condition from three independent experiments. (**C**) Representative confocal fluorescence micrographs of empty GUVs, CAHS12, CAHS12ΔC after incubation at 25 °C for 12 hours, 37 °C for 12 hours, 45 °C for 12 hours, and 60 °C for 2 hours. Elevated temperatures induced a marked reduction in vesicle size, consistent with heat-driven membrane remodeling or collapse.

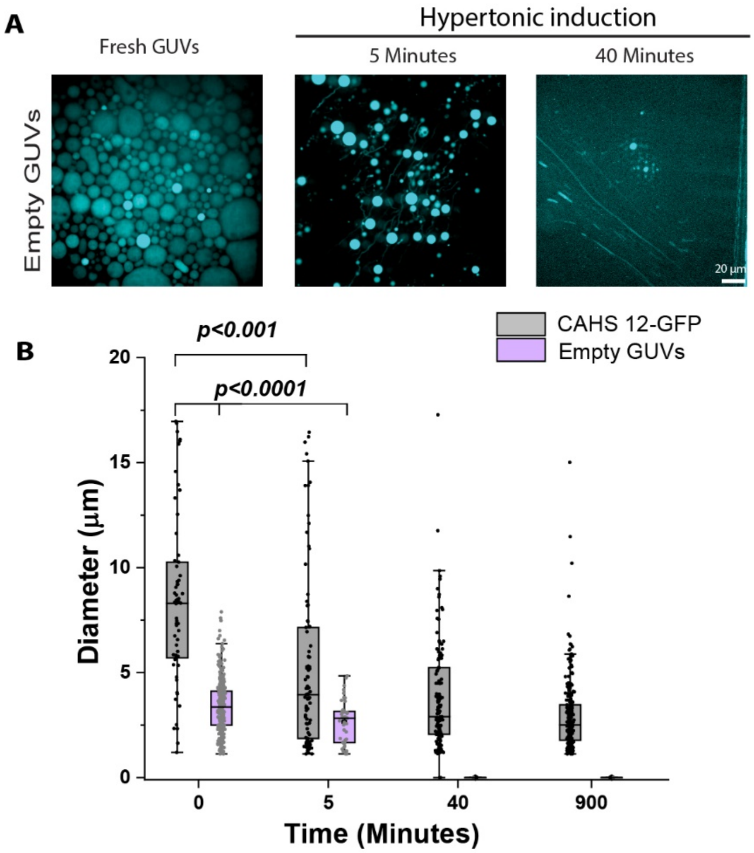


**Supplementary Fig. 23. Time-resolved morphological response of GUVs to hyperosmotic stress.** (**A**) After applying 2.5-fold hyperosmolar stress, empty GUVs were imaged using fluorescence microscopy after adding a hyperosmolar glucose solution. Scale bar, 20 µm. (**B**) Quantification of GUV diameter over time following the application of 2.5-fold hyperosmotic pressure. GUVs encapsulating CAHS12 or control solutions were imaged by confocal fluorescence microscopy at defined time points post-stress. CAHS12-stabilized vesicles exhibited slightly reduced shrinkage and preserved spherical morphology, whereas control GUVs rapidly collapsed or deformed. Data represent mean ± s.d. from >100 vesicles per condition, across three independent experiments.

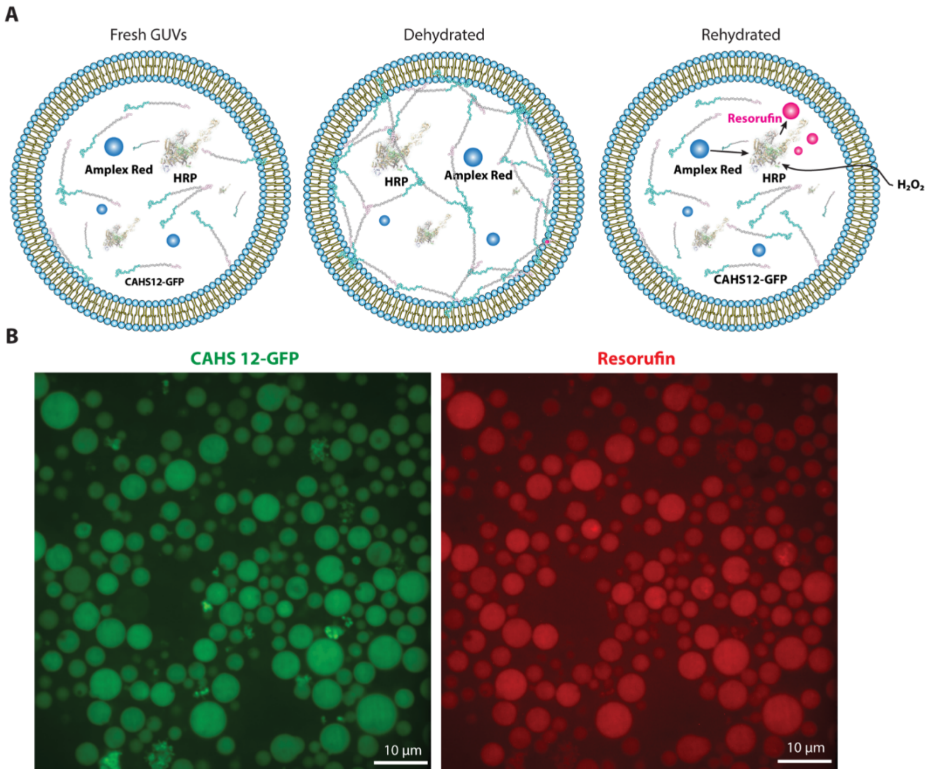


**Supplementary Fig. 24. Rehydration-activated enzyme catalysis within GUVs.** (**A**) Schematic illustration of the rehydration-triggered enzymatic reaction within GUVs. Upon rehydration, the encapsulated enzyme and non-fluorescent substrate are brought into proximity in the confined aqueous lumen, initiating the catalytic reaction that yields the fluorescent product, resorufin. (**B**) Fluorescence microscopy image showing the formation of fluorescent resorufin within individual GUVs following rehydration activation. Resolufin fluorescence indicates successful enzymatic conversion, demonstrating spatial confinement and functionality of the encapsulated catalytic system.

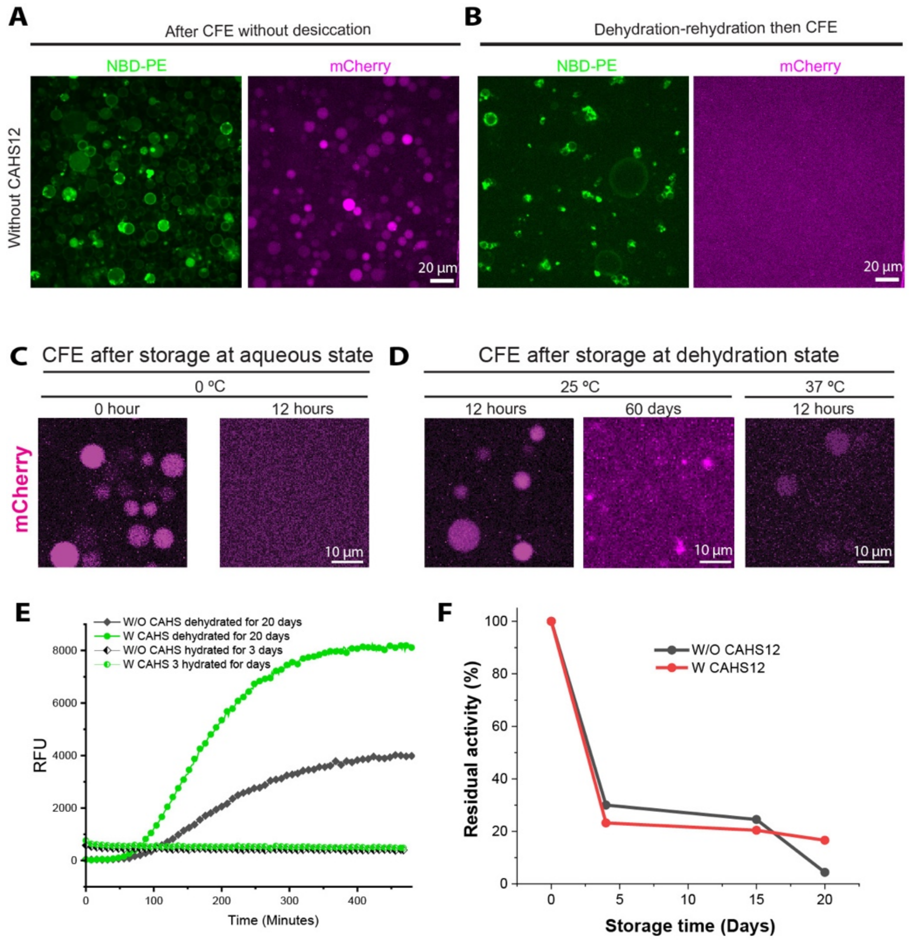


**Supplementary Fig. 25. Preservation of CFE activity in dehydrated GUVs under varying conditions.** Comparison of mCherry expression efficiency in CFE systems encapsulated within GUVs lacking CAHS12, before (**A**) and after dehydration and subsequent rehydration (**B**). Fluorescence intensity of mCherry was markedly reduced following the dehydration–rehydration cycle, indicating substantial loss of CFE activity in the absence of protective components. Quantification of CFE reaction efficiency in GUVs without (**C**) or with CAHS12-GFP (**D**) following dehydration at different storage temperatures. GUVs containing CAHS12-GFP exhibited enhanced preservation of CFE activity across temperature conditions, indicating a protective effect of the intrinsically disordered protein. (**E**) Relationship between residual water content and preserved CFE activity following dehydration. Higher water content was correlated with reduced preservation efficiency, underscoring the importance of controlled dehydration for long-term stability. Scale bars, 10 µm (C-D) and 20 µm (A-B). (**F**) Bulk (non-encapsulated) CFE reactions with or without CAHS12 were dehydrated and stored under identical conditions for different durations, followed by rehydration and assessment of translational activity via mCherry expression. Activity retention is reported relative to freshly prepared, non-dried bulk CFE controls.

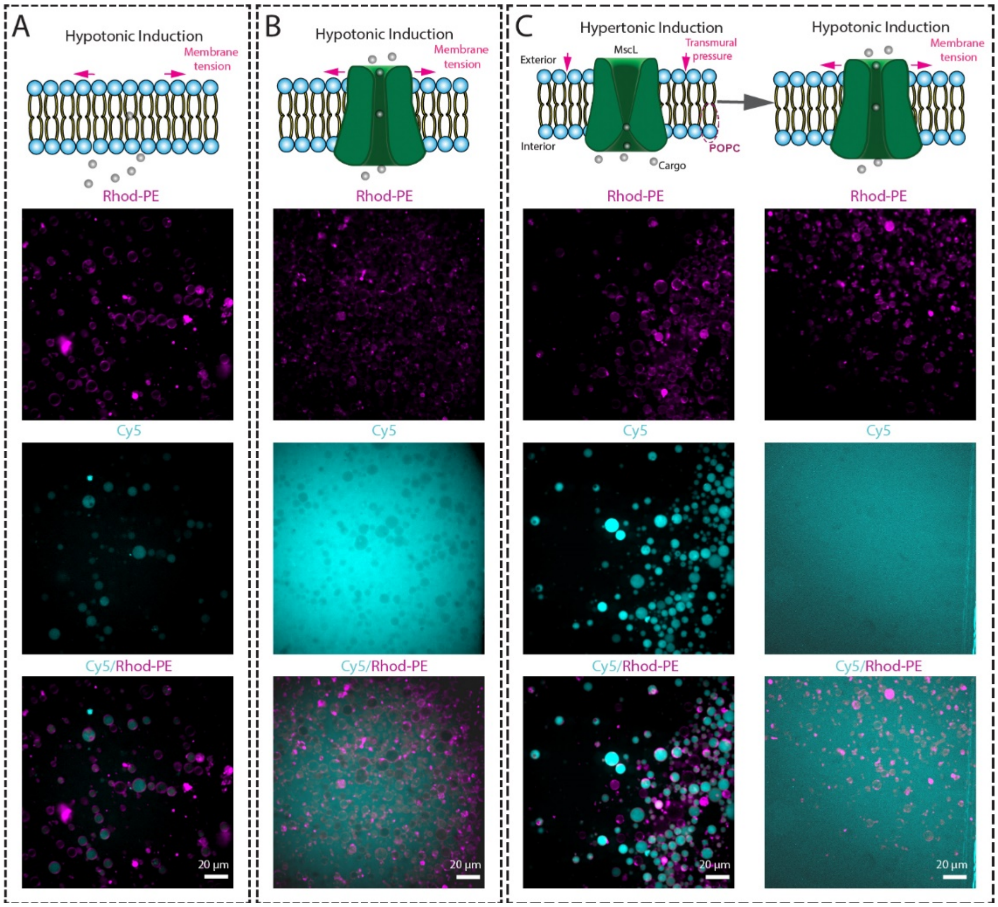


**Supplementary Fig. 26. Functional testing of MscL activity in GUVs after desiccation storage under osmotic stress.** (**A**) Fluorescence microscopy images of GUVs lacking MscL subjected to hypo-osmotic conditions following desiccation storage. The lipid membrane is labeled with Rhod-PE (magenta, top), while the internal cargo is Cy5 dye (cyan, middle); merged images (bottom) show retention of Cy5 fluorescence, indicating minimal leakage under membrane tension alone. (**B**) GUVs with MscL reconstituted in the membrane, subjected to hypo-osmotic induction, exhibit loss of encapsulated Cy5 fluorescence, indicating MscL activation and cargo release under membrane tension. (**C**) Sequential osmotic modulation experiment showing MscL-GUVs under hyperosmotic loading (cargo retained) followed by transfer to hypo-osmotic conditions, triggering MscL gating and Cy5 release. All images were acquired after a complete dehydration–rehydration cycle. Scale bars, 20 µm.

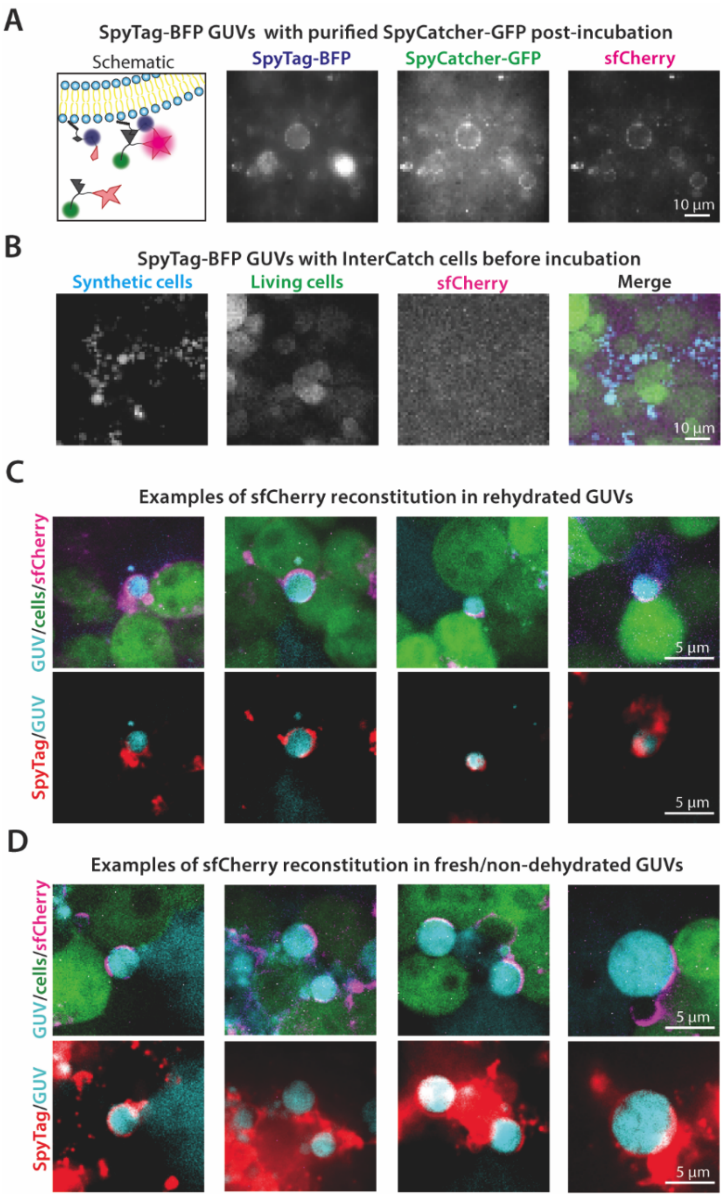


**Supplementary Fig. 27. Reconstitution of sfCherry between SpyTag-BFP–functionalized synthetic cells and InterCatch-expressing living cells.** (**A**) sfCherry reconstitution on SpyTag-BFP synthetic cells in the presence of purified SpyCatcher-GFP, a purified recombinant protein containing SpyCatcher, the sfCherry1–10 fragment, and GFP. (**B**) Representative image of SpyTag-BFP GUVs with InterCatch cells before incubation. (**C**) Four representative images of rehydrated, CAHS-encapsulating, SpyTag-BFP GUVs and InterCatch cells following a 4-hr incubation together. Note SpyTag-BFP signal is visible on the synthetic cell surface, with protein aggregation attributed to membrane clustering during GUV fabrication. (**D**) Four representative images of fresh CAHS-encapsulating, SpyTag-BFP GUVs and InterCatch cells following a 4-hr incubation together.

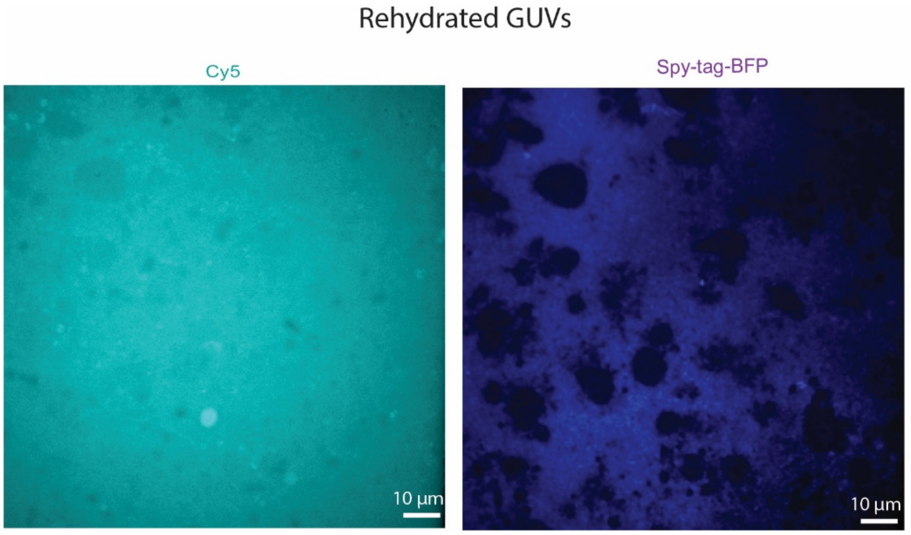


**Supplementary Fig. 28. CAHS12-free control for GUV stability after dehydration–rehydration.** Fluorescence microscopy images of GUVs lacking CAHS12 after dehydration and rehydration. GUV membranes are decorated with a SpyTag-BFP fusion protein covalently attached via a SpyCatcher lipid anchor, serving as a fluorescent marker for membrane-associated proteins (blue). Cy5 fluorescence reports soluble cargo. Following dehydration–rehydration, Cy5 signal is observed throughout the field, indicating cargo leakage from GUVs, while the SpyTag-BFP signal appears patchy and discontinuous, consistent with membrane disruption in the absence of CAHS12.

### 3. Supplementary Tables

**Supplementary Table 1** Sequence of CAHS12

| |
|---|
| MSHTHEQKFERVEERKVDDKKGLQEVRVGVDTGHGDPALNFTPTDATLVRTGGVGGTTAS<br>SHSSHMSSSTGGAVTGASQYSSTMHQEGGHMTTEASKNTSYTHTEVRAPVIDTAPPIISTGA<br>SGMAEQIVGQGFTASAARITGSSADVNIVETAEAREKRMRDEEKYAREQEAINRHADKDLEK<br>KTEAYRKEAEHEAEKIRKALEKQHERDIEFRKEVVGSTIEKQKAEIELEAKRAKAALEHERQLA<br>NDALERSKMHTDVQVTMDTAAGHTVSGGTTMSSSEQHSSSHSSSRTGGL |

**Supplementary Table 2** Sequence of CAHS12-GFP

| |
|---|
| MSHTHEQKFERVEERKVDDKKGLQEVRVGVDTGHGDPALNFTPTDATLVRTGGVGGTTAS<br>SHSSHMSSSTGGAVTGASQYSSTMHQEGGHMTTEASKNTSYTHTEVRAPVIDTAPPIISTGA<br>SGMAEQIVGQGFTASAARITGSSADVNIVETAEAREKRMRDEEKYAREQEAINRHADKDLEK<br>KTEAYRKEAEHEAEKIRKALEKQHERDIEFRKEVVGSTIEKQKAEIELEAKRAKAALEHERQLA<br>NDALERSKMHTDVQVTMDTAAGHTVSGGTTMSSSEQHSSSHSSSRTGGLGGGGSGGGGS<br>GGGGSMVSKGAELFTGIVPILIELNGDVNGHKFSVSGEGEGDATYGKLTLKFICTTGKLPVP<br>WPTLVTTLSYGVQCFSRYPDHMKQHDFFKSAMPEGYIQERTIFFEDDGNYKSRAEVKFEGD<br>TLVNRIELTGTDFKEDGNILGNKMEYNYNAHNVYIMTDKAKNGIKVNFKIRHNIEDGSVQLAD<br>HYQQNTPIGDGPVLLPDNHYLSTQSALSKDPNEKRDHMIYFGFVTAAAITHGMDELYK |

**Supplementary Table 3** Sequence of CAHS12-BFP

| |
|---|
| MSHTHEQKFERVEERKVDDKKGLQEVRVGVDTGHGDPALNFTPTDATLVRTGGVGGTTAS<br>SHSSHMSSSTGGAVTGASQYSSTMHQEGGHMTTEASKNTSYTHTEVRAPVIDTAPPIISTGA<br>SGMAEQIVGQGFTASAARITGSSADVNIVETAEAREKRMRDEEKYAREQEAINRHADKDLEK<br>KTEAYRKEAEHEAEKIRKALEKQHERDIEFRKEVVGSTIEKQKAEIELEAKRAKAALEHERQLA<br>NDALERSKMHTDVQVTMDTAAGHTVSGGTTMSSSEQHSSSHSSSRTGGLGGGGSGGGGS<br>GGGGSMSELIKENMHMKLYMEGTVDNHHFKCTSEGEGKPYEGTQTMRIKVVEGGPLPFAF<br>DILATSFLYGSKTFINHTQGIPDFFKQSFPEGFTWERVTTYEDGGVLTATQDTSLQDGCLIYN<br>VKIRGVNFTSNGPVMQKKTLGWEAFTETLYPADGGLEGRNDMALKLVGGSHLIANIKTTYRS<br>KKPAKNLKMPGVYYVDYRLERIKEANNETYVEQHEVAVARYCDLPSKLGHKLN |

**Supplementary Table 4** Sequence of CAHS12 ΔC

| |
|---|
| MSHTHEQKFERVEERKVDDKKGLQEVRVGVDTGHGDPALNFTPTDATLVRTGGVGGTTAS<br>SHSSHMSSSTGGAVTGASQYSSTMHQEGGHMTTEASKNTSYTHTEVRAPVIDTAPPIISTGA<br>SGMAEQIVGQGFTASAARITGSSADVNIVETAEAREKRMRDEEKYAREQEAINRHADKDLEK<br>KTEAYRKEAEHEAEKIRKALEKQHERDIEFRKEVVGSTIEKQKAEIELEAKRAKAALEHERQLA<br>NDALERSKMHTDVQVTMDT |

**Supplementary Table 5** Sequence of CAHS12 ΔN

| |
|---|
| AEAREKRMRDEEKYAREQEAINRHADKDLEKKTEAYRKEAEHEAEKIRKALEKQHERDIEFR<br>KEVVGSTIEKQKAEIELEAKRAKAALEHERQLANDALERSKMHTDVQVTMDTAAGHTVSGGT<br>TMSSSEQHSSSHSSSRTGGL |

**Supplementary Table 6** Sequence of CAHS12 ΔCΔN

| |
|---|
| AEAREKRMRDEEKYAREQEAINRHADKDLEKKTEAYRKEAEHEAEKIRKALEKQHERDIEFR<br>KEVVGSTIEKQKAEIELEAKRAKAALEHERQLANDALERSKMHTDVQVTMDT |

**Supplementary Table 7** Assignment of CAHS12 constructs to experimental assays

| **CAHS12 construct** | **Tag** | **Primary purpose** | **Assays used** |
|---|---|---|---|
| CAHS12 (full length) | None | Functional protection without tag interference | Long-term dry storage assays; interfacial tension measurements; thermal stability |
| CAHS12–GFP | GFP | Visualization of protein localization and assembly | Confocal imaging of GUVs; dehydration–rehydration dynamics; network formation; interfacial tension measurements; thermal stability |
| CAHS12–BFP | BFP | Visualization of protein localization and assembly | Confocal imaging of GUVs; Control to test tag effects on GUV protection |
| CAHS12 ΔN | None | Dissection of role of N-terminal region | GUV protection assays; interfacial tension measurements; stability comparisons |
| CAHS12 ΔC | None | Dissection of role of C-terminal region | GUV protection assays; interfacial tension measurements; stability comparisons |

10. WHAM – Grossfield Lab. http://membrane.urmc.rochester.edu/wordpress/?page_id=126.
